# Pipette: Encoding scientific literature into an executable Skill Graph for multi-agent bioinformatics

**DOI:** 10.64898/2026.04.08.717332

**Authors:** Chirag Gupta, Ananya Sharma

## Abstract

The cost of genomic sequencing has fallen by several orders of magnitude, yet data analysis remains a bottleneck concentrated among researchers with specialized computational expertise. While Large Language Models can generate bioinformatics code, they frequently produce incoherent multi-step workflows due to the absence of domain-specific analytical constraints. Here, we present Pipette, a multi-agent AI framework that orchestrates end-to-end bioinformatics workflows through natural language interaction, guided by a literature-derived Skill Graph. This directed, edge-weighted knowledge graph, extracted from over 20,000 peer-reviewed publications, constrains workflow generation to biologically valid analytical transitions, preventing incomplete or incoherent workflows. We benchmarked Pipette across four biological domains using published datasets: single-cell RNA-seq analysis of peripheral blood mononuclear cells and a human pancreas atlas, bulk RNA-seq differential expression in rice under environmental stress, and two structure-based computational drug design workflows. In ablation against two LLMs operating without Skill Graph constraints, Pipette matched or exceeded both baselines across all quantitative metrics while uniquely completing multi-step cross-domain transitions. We further evaluated Pipette on a clinical genomics task, where it executed an ACMG/AMP-compliant variant classification on a reference human genome. In all cases, Pipette recapitulated established biological and clinical findings while generating a fully reproducible, machine-readable provenance record. By reducing the computational expertise required to execute standard genomic analyses, Pipette lowers the barrier between sequencing data and biological insight for bench scientists. Pipette is available at https://pipette.bio.

## Introduction

Next-generation sequencing costs have fallen by several orders of magnitude over the past decade, and single-cell technologies now routinely profile tens of thousands of cells in a single experiment^1^. Yet the analytical capacity required to interpret this data has not scaled proportionally: the bottleneck in genomics has shifted decisively from data generation to data analysis^2^. A typical NGS experiment requires command-line proficiency, familiarity with multiple software tools, knowledge of appropriate statistical methods, and the ability to chain these into a reproducible workflow. For the majority of bench biologists, this creates a dependency on computational collaborators, slows research timelines, and introduces communication overhead that compounds with experimental complexity.

Large language models (LLMs) offer a potential route through this bottleneck. Beyond generating fluent text, recent LLMs exhibit agentic capabilities: the ability to plan multi-step actions, invoke external tools, evaluate intermediate outputs, and revise their approach accordingly^3,4^. This shift from passive text completion to goal-directed task execution has already attracted considerable interest in the life sciences^5–7^, with applications in single-cell genomics^8^, agriculture^9^, disease diagnosis^10^, and clinical decision support^11^. In chemistry, ChemCrow demonstrated that an LLM agent could autonomously plan and execute chemical syntheses by combining reasoning with domain-specific computational tools^12^, while Coscientist showed that such agents could independently design and carry out laboratory experiments with minimal human oversight^13^. These precedents suggest that agentic AI could similarly transform access to computational biology.

Several groups have begun addressing this opportunity directly. Biomni integrates 105 biomedical software tools and 59 databases into a general-purpose biomedical agent capable of tasks ranging from gene prioritization to protocol design^14^. AutoBA uses LLMs to propose analysis plans, generate code, and execute multi-omic workflows, though it requires local installation and computational infrastructure^15^. BIA demonstrated conversational automation of single-cell RNA-seq analysis including data retrieval and report generation, but remains limited to a single data modality^16^. BioAgents introduced a multi-agent architecture for troubleshooting bioinformatics pipelines using fine-tuned small language models with retrieval-augmented generation^17^. Other recent systems include CompBioAgent for single-cell data exploration^18^, CellAtria for dialogue-driven scRNA-seq standardization^8^, and Talk2Biomodels for interacting with kinetic models of biological systems^19^. Each addresses a component of the problem, but none provides a unified framework that spans multiple sequencing modalities while maintaining the transparency of agent reasoning and the reproducibility of data provenance that deployment in a research laboratory demands.

A core limitation shared by these systems is susceptibility to workflow incoherence during multi-step planning^20^. LLMs are trained on vast general corpora, but they lack explicit representations of the domain-specific constraints that govern bioinformatics: which analytical operations are compatible, what data types flow between tools, and which procedural transitions are sanctioned by community practice. Equipping agents with curated standard operating procedures – individual “Skills” documents that standardize the execution of discrete computational tasks – partially addresses this. However, a flat library of skills is insufficient for workflow orchestration. The relationships between tools, data types, and analytical stages are distributed across thousands of published methods sections, and an unconstrained LLM can correctly execute one skill yet hallucinate an incompatible downstream transition. What is needed is not a larger skill library, but a topological map of valid analytical pathways: a structured knowledge representation where nodes encode discrete skills and directed, weighted edges enforce literature-backed, data-type-compatible transitions.

Here, we present Pipette, a multi-agent AI framework that translates natural language queries into rigorous, reproducible bioinformatics workflows. At the core of Pipette is the Skill Graph – a directed, weighted network of analytical transitions computationally extracted from over 20,000 full-text peer-reviewed publications in PubMed Central using fine-tuned biomedical language models. By grounding workflow planning in empirically validated, data-compatible pathways, the Skill Graph constrains the agent’s reasoning to transitions supported by community consensus, reducing incoherent or fabricated analytical steps, while reducing token usage. Pipette operates through a multi-stage architecture in which natural language queries are decomposed into structured computational tasks, domain-specific code is generated and executed, methodology is subjected to an independent LLM-driven audit, and outputs are synthesized into interpretable biological results with a complete, machine-readable provenance record. We benchmarked Pipette across four technically demanding domains, including single-cell transcriptomic clustering, multifactorial bulk RNA-seq analysis, structure-based drug design, and ACMG-compliant clinical variant classification. We show that in each case, it recapitulated established biological and clinical findings while maintaining full analytical reproducibility.

### Architecture of the Pipette Multi-Agent Framework

Pipette is a multi-agent framework that executes bioinformatics workflows from natural language input (**Figure 1**). The platform runs on scalable cloud infrastructure and is organized into mainly two functional layers: a conversational interface and an orchestration engine. The conversational layer is governed by a Copilot Agent, which interprets natural language queries, classifies user intent (e.g., explain, execute, help), resolves biological ambiguities through clarifying dialogue, and routes structured analytical tasks to the appropriate compute queue with the required resources. Once a task is routed, an Orchestrator Agent dispatches it through a six-stage analytical pipeline, supported by an elastic compute cluster that dynamically allocates resources alongside ephemeral and persistent object storage.

**Figure 1.**
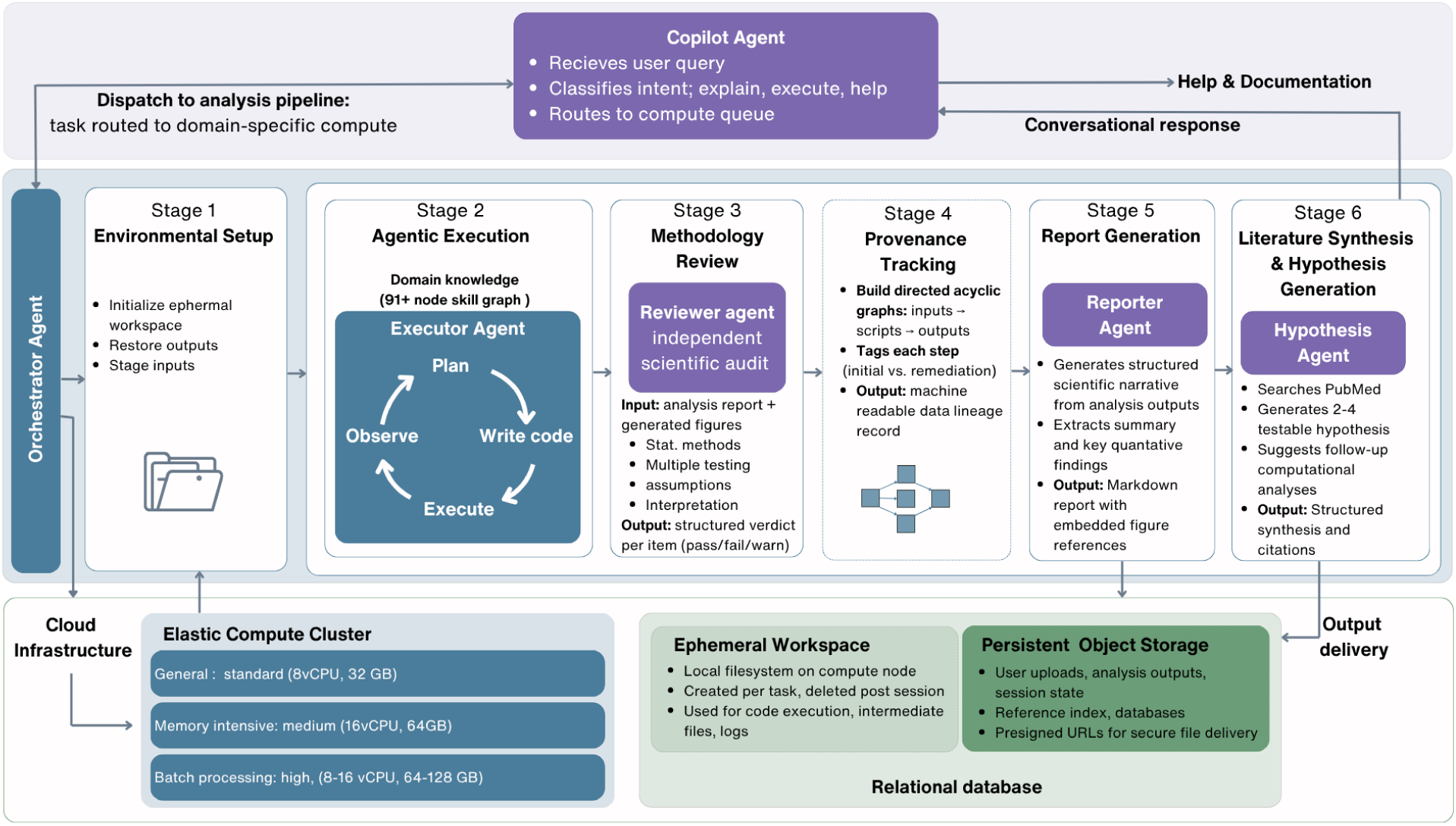
Architecture of Pipette.bio, a multi-agent AI framework for autonomous bioinformatics analysis. The system comprises three layers. **(Top)** A conversational interface where the Copilot Agent classifies user intent – responding directly to conceptual questions or dispatching analysis requests to the pipeline with domain-aware queue routing. **(Middle)** A six-stage orchestrated analysis pipeline coordinated by a deterministic orchestrator. Stage 1 initializes an ephemeral workspace and restores outputs from prior conversation turns for multi-turn continuity. Stage 2 houses the Executor Agent, which operates in an iterative tool-use loop (Plan, Write Code, Execute, Observe) across bash, Python, and R, with access to a library of 91+ injectable domain knowledge modules (skills) covering RNA-seq alignment, differential expression, variant calling, single-cell analysis, metagenomics, drug design, and other domains. Stage 3 employs an independent Reviewer Agent that audits the analysis for statistical rigor, assumption validity, and methodological correctness, issuing structured pass/warn/fail verdicts; methodological failures trigger a remediation loop (red dashed arrow) that returns to the Executor for targeted correction. Stage 4 performs deterministic provenance tracking, constructing a directed acyclic graph linking inputs, scripts, and outputs for full reproducibility. Stage 5 uses a Reporter Agent to generate a structured scientific narrative with key quantitative findings. Stage 6 deploys a Hypothesis Agent that searches biomedical literature databases and generates testable mechanistic hypotheses grounded in the user’s results. Finally, the agent packages outputs into a canonical directory structure and uploads them to persistent storage with time-limited secure access URLs. (Bottom) Cloud infrastructure comprising five domain-specific elastic compute queues that scale to zero when idle – ranging from standard (4 vCPU, 16 GB) for general analysis to high-memory (32 vCPU, 128 GB, 500 GB disk) for bulk RNA-seq alignment, alongside an ephemeral workspace (task-scoped local filesystem, deleted after finalization), persistent object storage (user uploads, analysis outputs, session state, and several pre-built reference indexes), and a relational database storing users, sessions, and provenance audit trails.

The pipeline begins with Stage 1 (Environmental Setup), which initializes an isolated ephemeral workspace and stages the necessary inputs. The core computation occurs in Stage 2 (Agentic Execution), where an Executor Agent draws on domain knowledge from the Skill Graph (described in the next section) to navigate an iterative plan, write code, execute, observe loop. This cycle enables the agent to dynamically load tools, configure organism-specific parameters, and diagnose and recover from intermediate execution errors.

To enforce methodological standards, Stage 3 (Methodology Review) introduces an independent Reviewer Agent that audits the executor’s code and generated figures, evaluating statistical methods, multiple testing corrections, and analytical assumptions. An iterative reviewer-executor loop ensures that analyses proceed only after the Reviewer Agent issues a structured pass verdict.

Stage 4 (Provenance Tracking) captures a machine-readable data lineage for each workflow. Directed acyclic graphs (DAGs) map inputs and scripts to final outputs, recording exact software versions, parameters, random seeds, and intermediate states at each step.

In the final stages, Pipette translates analytical outputs into interpretable biological results. Stage 5 (Report Generation) uses a Reporter Agent to extract key quantitative findings and compile a structured Markdown report with embedded figure references. Stage 6 (Literature Synthesis and Hypothesis Generation) employs a Hypothesis Agent that queries external knowledge bases, including PubMed, to contextualize findings against current literature, generate testable hypotheses, and suggest follow-up computational analyses based on the Skill Graph transitions. The complete output, comprising the scientific narrative, provenance record, and hypotheses, is returned to the user through the conversational interface.

### The Skill Graph

We define a skill as a self-contained analytical operation ( e.g., read alignment, pathway enrichment etc.) specified by its trigger conditions, required inputs, executable code, and typed outputs, which the Executor Agent loads and executes at runtime. To constrain workflow planning to biologically valid analytical sequences, we construct the Skill Graph: a directed, weighted network of 91 analytical skills spanning nine biological domains (**Figure 2**). Each node represents a discrete analytical skill, linked by directed edges that encode valid sequential transitions between skills, weighted by literature co-occurrence frequency. The Skill Graph is generated through a dual-track extraction pipeline applied to approximately 20,000 PubMed Open Access papers and 800 manually curated notes **(Figure 2A**).

**Figure 2.**
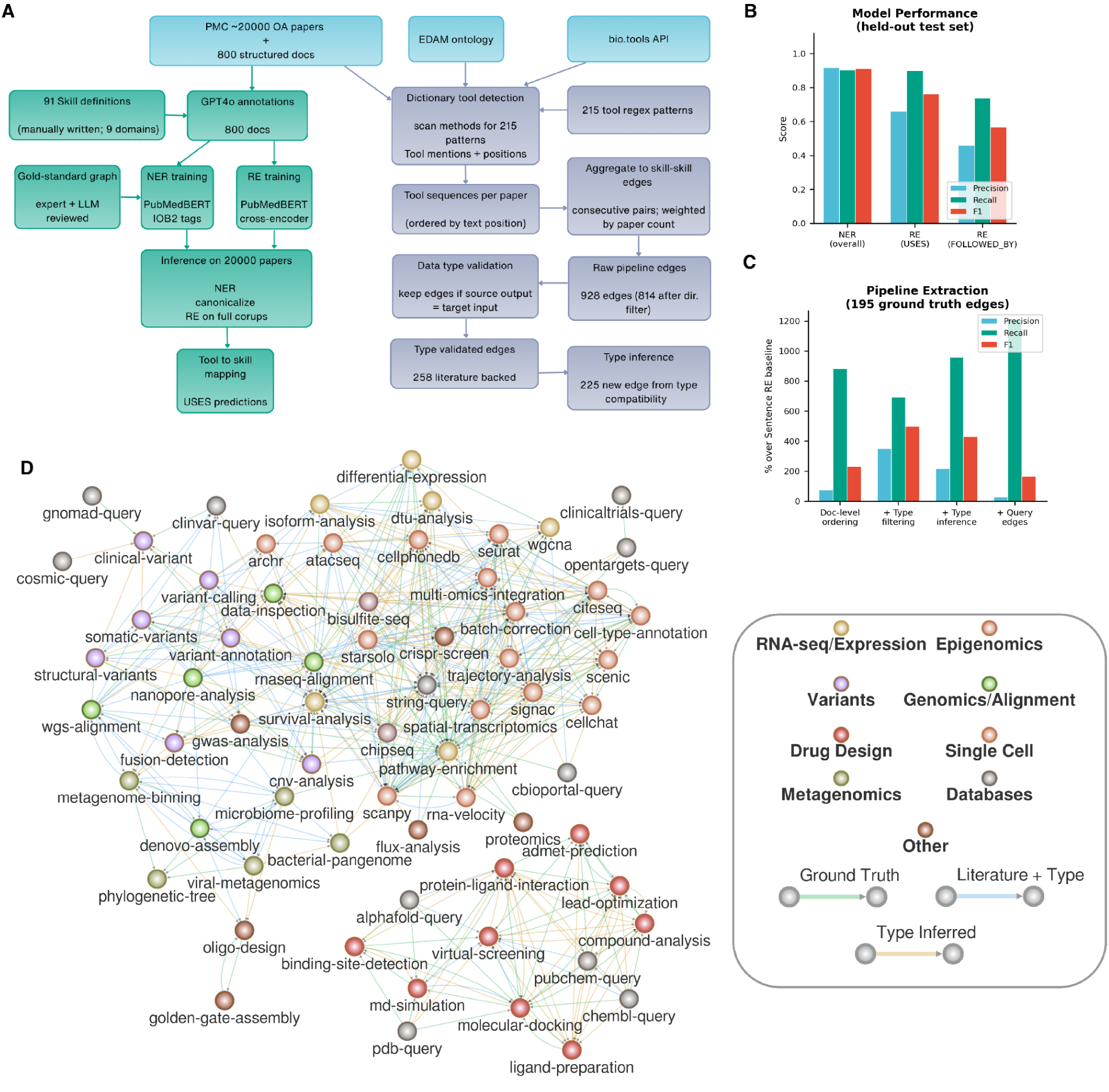
Skill Graph: automated construction of a bioinformatics workflow knowledge graph from scientific literature. **A)** Overview of the Skill Graph construction pipeline. Starting from ∼20,000 open-access PMC papers and 800 structured notes across 9 domains, Named Entity Recognition (NER) and Relation Extraction (RE) models were trained on PubMedBERT with IOB2 tags and cross-entropy loss, respectively. Dictionary-based tool detection using EDAM ontology and bio.tools API identified 215 tool regex patterns. Tool mentions and positions were aggregated into skill-skill edges weighted by paper count, yielding 628 raw pipeline edges. Data type validation retained edges where source outputs matched target inputs, producing 258 literature-backed edges. An additional 225 edges were inferred from type compatibility alone. **B)** Model performance on held-out test set. Grouped bar chart showing Precision (blue), Recall (teal), and F1 (red) for NER (overall entity recognition), RE (USES relation type), and RE (FOLLOWED_BY relation type). **C)** Pipeline extraction accuracy expressed as percentage improvement over the Sentence RE baseline, evaluated against 195 expert-curated ground truth edges. Four successive enrichment stages are shown: document-level ordering, type filtering, type inference, and query edge addition. The Y-axis is on a symmetric log scale. **(D)** The resulting SkillGraph comprising 78 bioinformatics skills and 483 directed edges. Nodes are colored by domain: Genomics/Alignment (olive), Variants (clay), RNA-seq/Expression (sand), Single Cell (terracotta), Epigenomics (dusty rose), Drug Design (burnt sienna), Metagenomics (dark khaki), Databases (warm gray), and Other (brown). Node size scales with the number of associated tools. Edges are colored by evidence type: green (ground truth, confirmed in expert-curated graph), blue (literature + type, co-occurrence in papers with data type compatibility), and gold (type inferred, no direct paper co-occurrence but compatible input/output types).

The first track focuses on mapping specific software tools to their corresponding conceptual skills using machine learning. We utilized GPT-4o to generate joint Named Entity Recognition (NER) and Relation Extraction (RE) annotations across an 800-document seed corpus. We then fine-tune PubMedBERT-based models on these annotations. The resulting NER model achieves high accuracy in identifying tools, operations, and data types (overall F1 = 0.91), while the RE model reliably identifies tool-to-operation “USES” relationships (F1 = 0.76) **(Figure 2B**). Running inference across the full 20,000-paper corpus successfully maps real-world bioinformatics tools to 78 of the 91 predefined skills.

The second track extracts pipeline structure by identifying valid sequential transitions between skills. We define a dictionary of 215 regex patterns linked to EDAM ontology^21^ and the bio.tools API^22^. We scan the methods sections of papers, and record the sequential text positions of the matched tools to extract candidate workflow orderings weighted by paper co-occurrence. A key finding during this phase is that document-level sequential ordering substantially outperforms sentence-level relation extraction for pipeline discovery (**Figure 2C**). Standard sentence-level RE matches only 6.2% (12 of 195) of expert-curated ground-truth edges, as analytical steps in methods sections often span multiple paragraphs. Document-level ordering recovers 60.5% (118 of 195) of ground-truth transitions.

To eliminate logically invalid transitions, we apply data type validation: a candidate edge from an upstream to a downstream skill is retained only if the output data types of the former are compatible with the input data types of the later. This filtering improves pipeline extraction precision from 14.5% to 37.2% and enables inference of new, logically valid edges between skills that are never explicitly co-observed in the literature. The two tracks then converge to form the final Skill Graph. The ML track populates nodes with curated toolsets, while the document-extraction track, enriched by type inference, establishes the connective topology. The resulting graph comprises 258 literature-derived edges and over 200 type-inferred edges, recovering 65.1% of all expert-curated ground-truth transitions.

These results can be contextualized against existing biomedical information extraction approaches. Standard sentence-level RE, the dominant paradigm for biomedical relation extraction^23,24^, achieves only 6.2% recall on our pipeline extraction task, consistent with the known difficulty of cross-sentence relation extraction in scientific text^25^. Sebe et al. (2025), the most directly comparable prior work, reported 70.4 F1 for bioinformatics tool NER using SciBERT but did not evaluate pipeline edge recovery^23^. Our PubMedBERT-based NER achieves higher F1 (0.91 vs. 0.70) on a broader entity vocabulary (TOOL, OPERATION, DATA_TYPE vs. tool mentions alone), likely reflecting both the larger training corpus (800 vs. ∼200 documents) and the inclusion of internal lab notes that provide explicit tool-to-operation descriptions absent from research papers.

The SkillGraph is hosted at skillgraph.pipette.bio, where users can navigate the graph interactively and extract relevant workflows for their own agentic systems.

### Pipette’s behavioral metrics and system validation

To characterize Pipette’s operational behaviour in production, we analysed pipeline executions across four computational domains over a two-month deployment period. Bulk NGS and downstream analyses account for the majority of queries (67.4%), followed by single-cell (24.4%), microbiome (6.4%), and drug design (1.8%) (**Figure 3A**). The agent invoked 48 Skill Graph nodes, with differential expression, scanpy-based single-cell analysis, and bulk RNA-seq alignment as the most frequently used skills (**Figure**. **3B**). During multi-step analyses, the agent traversed multiple distinct Skill Graph edges, chaining sequential analytical skills in patterns consistent with established bioinformatics workflows. The most common is cBioPortal query followed by survival analysis, RNA-seq alignment followed by differential expression, and differential expression followed by pathway enrichment (**Figure 3C**). Cross-domain transitions, including a drug design chain (PubChem query → ADMET prediction → AlphaFold query) and a clinical genomics path (ClinVar query → gnomAD query), demonstrate multi-step reasoning beyond single-domain analyses.

**Figure 3.**
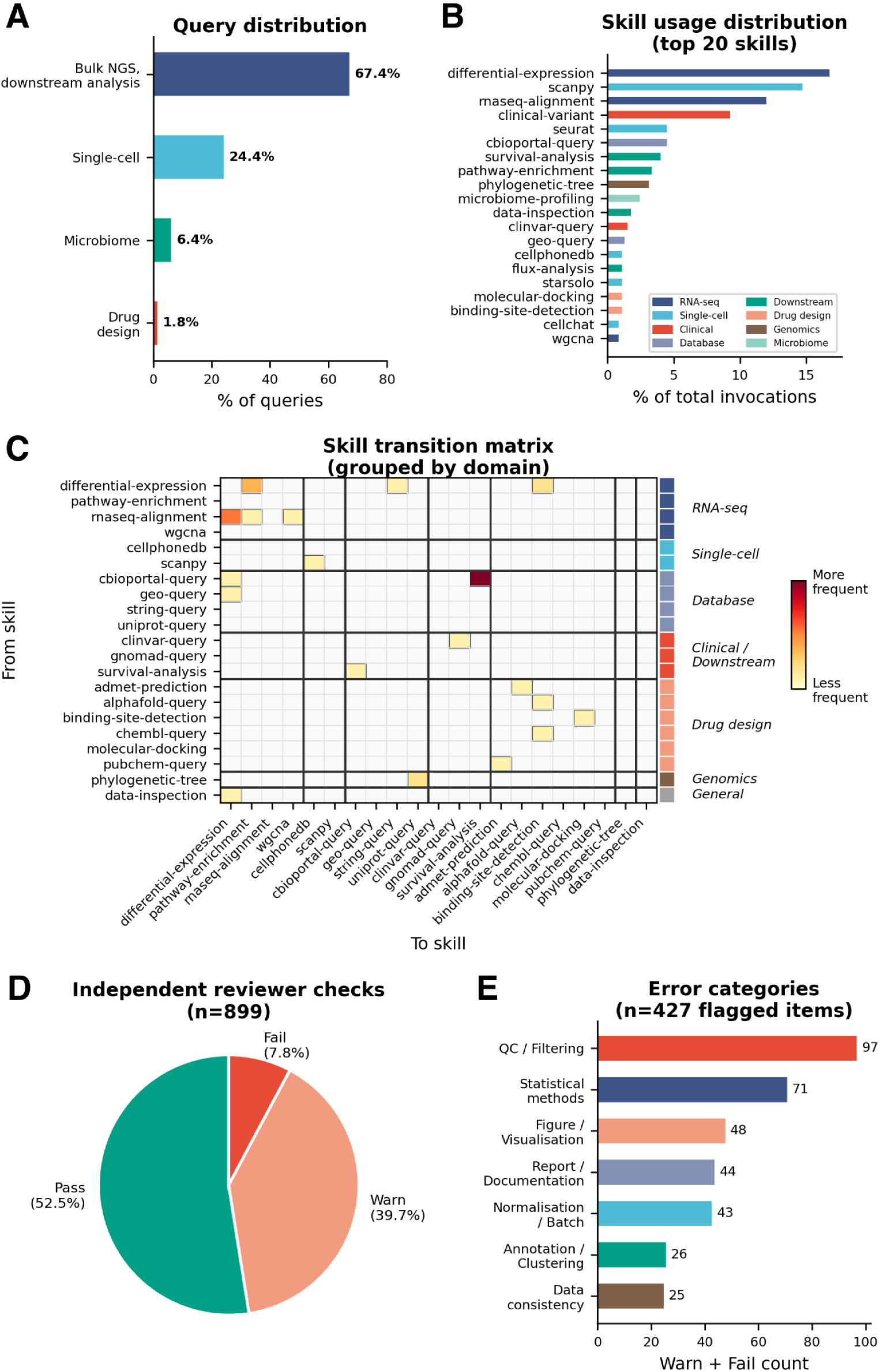
Pipette behavioural metrics and system validation across production deployments. **A)** Distribution of autonomous pipeline queries across four computational domains during a two-month deployment period. Percentages reflect proportion of total query volume. **B)** Skill usage distribution showing the 20 most frequently invoked skills (of 48 active), coloured by analytical domain. Differential expression, single-cell analysis (Scanpy, Seurat), and RNA-seq alignment together account for over 45% of all skill invocations. **C)** Skill Graph transition matrix showing directed skill-to-skill transitions observed during production analyses, grouped by analytical domain. Cell values indicate transition count; domain colour blocks on the right axis indicate skill category. Thick lines separate domain groups. The dominant transitions – cBioPortal → survival analysis, RNA-seq alignment → differential expression — correspond to canonical bioinformatics workflows. **D)** Independent reviewer check outcomes across 899 quality-check items from 125 analysis sessions. Pass: methodology meets standards; Warn: suboptimal but non-critical; Fail: remediation required before final report. **E)** Error categories flagged by the Reviewer Agent (warn + fail verdicts, n=427 items). QC/filtering and statistical method concerns dominate, mirroring the error classes most commonly raised in human peer review of bioinformatics manuscripts.

Further, aggregating 125 reports comprising 899 individual quality-check items by Pipette’s Reviewer Agent, we found that 52.5% of checks passed, 39.7% received warnings indicating suboptimal but non-critical methodology, and 7.8% were flagged as failures requiring remediation (**Figure 3D**). The most frequently flagged error categories are QC and filtering issues (n=97), such as missing doublet removal or inadequate gene-count thresholds, and statistical method concerns (n=71), predominantly absent or unspecified multiple testing correction (**Figure 3E**). Figure and visualisation errors (n=48), report documentation gaps (n=44), and normalisation or batch effect concerns (n=43) were also common. These results demonstrate that the Reviewer Agent functions as a substantive quality gate, systematically catching the same classes of methodological errors – QC thresholds, statistical rigour, and figure-text consistency – that human peer reviewers routinely flag in bioinformatics manuscripts.

### Benchmarking Pipette with Real-World Datasets

To evaluate Pipette under realistic conditions, we select six datasets that span different organisms, data modalities, and biological questions. For each case study, we use only the platform’s conversational interface to drive the analysis from raw data to biological interpretation. No code was written manually at any stage. The agent’s output files, generated codes, reasoning, and provenance records are provided as supplementary data on the project’s GitHub repository (see Methods). We compare these results against published analyses of the same datasets where possible to assess whether the platform reproduces established biological findings.

#### Single-cell RNA-seq

To evaluate Pipette’s capacity to autonomously execute single-cell workflows, we tasked the agent with the end-to-end analysis of a large-scale single-cell RNA-sequencing (scRNA-seq) dataset. Specifically, we provided Pipette with the raw gene-barcode matrices (matrix.mtx, barcodes.tsv, genes.tsv) from the landmark 68,000 peripheral blood mononuclear cell (PBMC)^1^ and instructed to perform a standard clustering and annotation analysis (see Methods).

Acting entirely autonomously, Pipette successfully formulated and executed a best-practice scRNA-seq pipeline. The agent initially applied standard quality control (QC) filters, retaining cells expressing between 200 and 3,000 genes, and restricting mitochondrial read content to below 20%. Following this stringent filtering, Pipette retained 68,548 of the original 68,579 cells (99.95% retention). The agent then normalized the data to 10,000 counts per cell, identified the top 2,000 highly variable genes, performed principal component analysis (PCA), and clustered the dataset using the Leiden algorithm at a resolution of 0.5.

For biological interpretation, Pipette seamlessly integrated automated cell type annotation by applying the CellTypist^26^. Utilizing the ‘Immune_All_Low.pkl’ model and employing a robust majority-vote scheme per cluster, the pipeline resolved 16 distinct Leiden clusters mapped to 10 broad immune cell types. The agent successfully generated a two-dimensional UMAP embedding reflecting these comprehensive annotations, clearly delineating major lymphoid and myeloid lineages including distinct T cell subsets, B cells, monocytes, and NK cells **(Figure 4A).** We next evaluated the accuracy of the AI agent’s autonomous workflow by comparing its derived cell type compositions against the manually annotated results published in the original study^1^. We observed that Pipette’s automated annotations exhibited striking concordance with the established reference baseline. A linear comparison of broad cell type proportions yielded a highly significant Pearson correlation of r=0.959 ( p = 1.63 × 10-4) **(Supplementary figure S1A)**.

**Figure 4:**
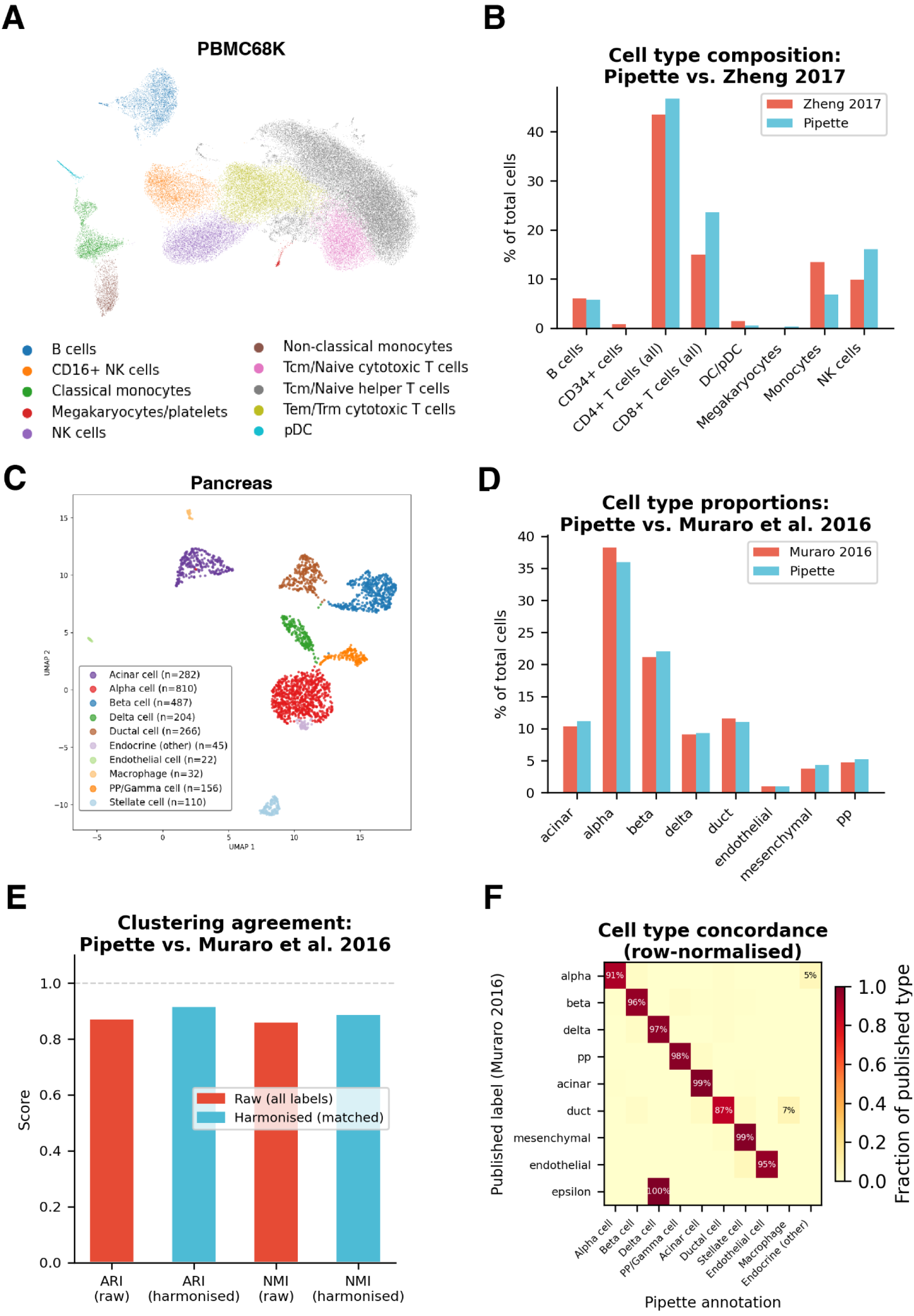
Evaluation of Pipette to execute scRNA-seq workflows using the PBMC 68K dataset. **A)** UMAP plot generated by Pipette showing cell clusters uniquely colored according to the annotations from CellTypist. **B)** Grouped bar chart comparing the percentage of total cells assigned to each major immune lineage by Pipette (CellTypist, blue) and as reported in the original reference publication (Zheng et al. 2017, red). Cell types are derived from Leiden clustering (resolution 0.5) followed by CellTypist ‘Immune_All_Low’ majority-vote annotation or reference-based classification using purified PBMC subpopulations (Zheng 2017). Note that Zheng 2017 does not report megakaryocytes as a separate category and leaves approximately 10% of cells unclassified, which contributes to apparent differences in CD8+ T cell and monocyte proportions. **C-F) Concordance of autonomous single-cell analysis with published cell type annotations in human pancreas. C)** UMAP of pancreas cell clusters and annotations generated by Pipette. **D)** Cell type proportions compared between the AI agent and the published reference across seven shared pancreatic cell types. Agent labels were harmonised to published nomenclature. **E)** Adjusted Rand Index (ARI) and Normalised Mutual Information (NMI), shown for both raw labels and harmonised labels (after excluding macrophage and mixed endocrine clusters without published equivalents). Harmonised ARI = 0.92, NMI = 0.89. **F)** Row-normalised confusion heatmap comparing published per-cell labels (rows) against agent annotations (columns). Diagonal dominance confirms strong agreement across all major cell types, with endothelial cells matched at 95% accuracy. Off-diagonal entries reflect reassignment of a subset of published beta cells to an Endocrine (other) cluster expressing mixed endocrine markers.

Detailed compositional analysis reveals minor discrepancies in specific lineage proportions between the agent and the reference **(Figure 4B).** Notably, Pipette appears to slightly overestimate the proportions of CD4+ T cells, CD8+ T cells, and NK cells, while underestimating monocytes relative to the original publication **(Supplementary figure S1B)**. However, these apparent differences are largely a computational artifact of differing annotation thresholds. In the original study, the reported group proportions sum to only ∼90%, as approximately 10% of cells were excluded from reported totals due to low-confidence classifications^1^. In contrast, Pipette’s automated CellTypist pipeline forces a classification for the entire post-QC dataset. Adjusting for this 10% unannotated baseline offset would structurally reduce the observed overestimates in the CD8+ and NK compartments.

Further, to quantitatively assess Pipette’s cell type annotation accuracy against another independent ground truth, we analysed the human pancreas CEL-seq2 dataset^27^(GSE85241; 3,072 cells, 4 donors), for which per-cell type labels are publicly available. Starting from the raw count matrix, the agent autonomously performed MAD-based QC filtering (retaining 2,414 cells, normalization, HVG selection with donor-aware batch handling, PCA, Leiden clustering (resolution 0.5, yielding 10 clusters), and cell type annotation using an overlap-score method against a curated pancreatic marker dictionary **(Figure 4C)**.

The agent correctly identified all seven major pancreatic cell types, including alpha, beta, delta, gamma/PP, acinar, ductal, stellate, endothelial cells (0.9%, matched at 95% accuracy against published labels), and macrophages (1.3%), with proportional concordance of r = 0.998 (p = 2.55 × 10⁻⁸) across eight shared cell types against the published reference **(Figure 4D, Supplementary figure S1C).** Pipette recovered >50% of published marker genes **(Supplementary figure S1D).** Harmonised ARI of 0.92 and NMI of 0.89 indicate strong clustering agreement when excluding macrophage and mixed endocrine clusters without published equivalents; the raw ARI of 0.87 reflects the agent’s additional resolution of populations not annotated in the original publication **(Figure 4E)**. Per-cell comparison against the published labels confirmed strong diagonal dominance in the row-normalised confusion heatmap, with all major cell types exceeding 87% concordance **(Figure 4F).** Only epsilon cells (n = 3) were not resolved as an independent cluster, reflecting insufficient cell numbers rather than analytical error.

Overall, this validation confirms that Pipette can ingest raw transcriptomic data and reliably execute sophisticated, reproducible, and literature-concordant analytical workflows.

#### Bulk RNA-seq

To further evaluate Pipette’s capacity for autonomous statistical modeling and handling of complex experimental designs, we tested its ability to perform differential gene expression analysis from a raw bulk RNA-seq count matrix. We provided Pipette with an 80-sample rice leaf count matrix (GEO accession GSE295637) from a recent study^28^ and instructed to identify differentially expressed genes (DEGs) across four specific stress contrasts.

Pipette correctly applied an appropriate pre-filtering threshold (≥ 10 counts in ≥ 4 samples), retaining 24,338 genes, which is highly consistent with the 24,326 tested genes reported in the original study. To evaluate differential expression, the agent autonomously formulated a DESeq2 negative-binomial generalized linear model (GLM)^29^. Importantly, Pipette correctly parsed the experimental metadata to include both sequencing batch and leaf segment as additive covariates alongside the primary biological condition (∼ batch + segment + condition). Moreover, Pipette perfectly recapitulated the core biological findings of the original study. Pipette-generated DEG counts mirror the published patterns: combined heat and drought stress (D40 vs W30) elicits the most profound overall transcriptional response, heat shock alone (W40 vs W30) is a considerably stronger stimulus than isolated drought (D30 vs W30), and transcriptomic downregulation clearly dominates the cellular response to heat stress.

However, quantitative comparisons reveal a systematic over-inflation in the absolute number of DEGs identified by Pipette. When matched to the strict significance threshold utilized by the original study (FDR < 0.01 and log 2 FC ≥1), Pipette identifies between 11.2% and 24.4% more DEGs across the four primary contrasts **(Figure 5A**). For example, in the D30 vs W30 contrast, the agent identifies 2,805 DEGs compared to the published 2,254 (+24.4%), and in the D40 vs W30 contrast, it identifies 6,577 compared to 5,539 (+18.8%). As Pipette utilized the raw count matrix directly, this residual discrepancy likely reflects minor differences in computational environments, such as DESeq2 package versions, exact implementation of independent filtering, or the potential absence of log fold-change (LFC) shrinkage (e.g., apeglm) in the agent’s default pipeline, which can systematically alter test statistics for low-count genes.

**Figure 5:**
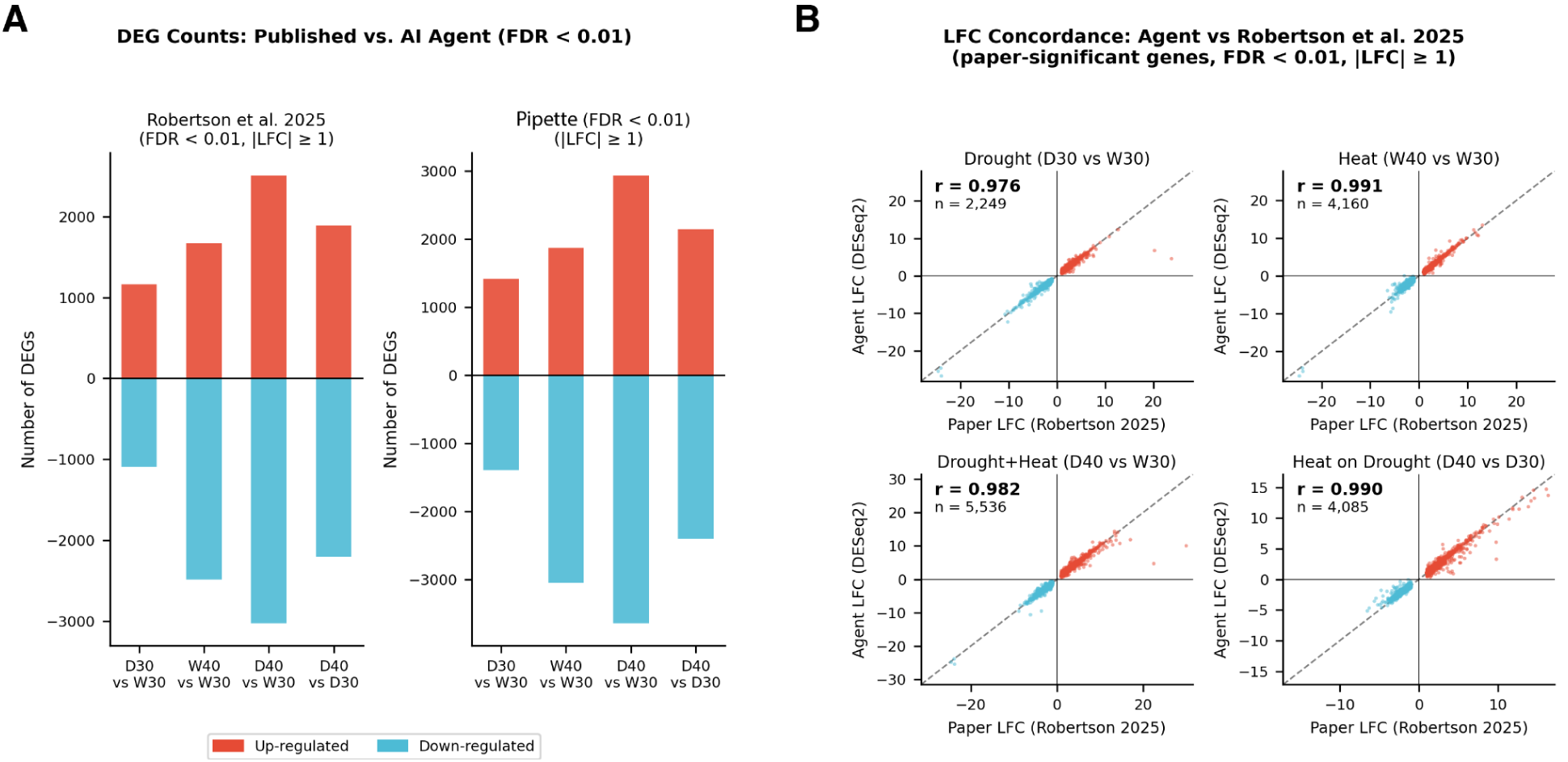
Evaluation of Pipette’s differential expression analysis workflows. **A)** Diverging bar charts comparing DEG counts (up = positive, down = negative) at matched FDR < 0.01 between Robertson et al. 2025 (left) and the AI agent recalculated at the same threshold (right). The agent consistently over-estimates DEG counts by ∼11–24%, attributable to the absence of LFC shrinkage. Colours: red = up-regulated, blue = down-regulated. **B)** Four-panel scatter plot showing gene-level log₂FC concordance between the AI agent’s DESeq2 output and Robertson et al. 2025 (Table S2) across all four contrasts. Only paper-significant genes (FDR < 0.01, |LFC| ≥ 1) are shown. Points coloured red (up-regulated in paper) or blue (down-regulated). Dashed line = identity (y = x). Pearson r ≥ 0.976 for all contrasts, confirming near-identical fold-change estimates.

Despite these minor thresholding and methodological differences, the underlying gene-level effect sizes computed by Pipette demonstrates near-perfect concordance with the originally published results (**Figure 5B**, right). When comparing the LFCs of paper-significant genes, the Pearson correlations between the Pipette and the original study were exceptionally high across all experimental conditions. The correlations ranged from r=0.976 (p<0.0001, n=2,249) for the isolated drought contrast to r = 0.991 (p<0.0001, n=4,160) for the heat shock contrast.

This high fidelity demonstrates Pipette’s ability to autonomously parameterize and execute robust, multivariate statistical analyses on raw transcriptomic data, yielding effect size estimates that are essentially at par with human-conducted analysis.

#### Drug Designing task-1: Imatini – ABL1 molecular docking

To evaluate Pipette’s proficiency in complex structure-based drug design, we performed two tasks. In the first, we tasked the agent with predicting the binding mode of the chemotherapeutic imatinib within the ATP-binding cleft of human ABL1 kinase (PDB: 2HYY). This task tested the agent’s capacity for autonomous methodological oversight and dynamic error recovery. During an initial execution pass, the independent Reviewer Agent flagged two critical chemoinformatic omissions: the absence of explicit physiological pH (7.4) protonation states and the lack of a quantitative crystal-pose validation. In response, the Executor Agent dynamically self-corrected the pipeline. To achieve the correct physiological state, the agent performed stringent pKa analysis on imatinib’s piperazinyl nitrogens (identifying the neutral form as the dominant species at pH 7.4) and attempted to prepare the receptor using PDB2PQR. When the agent detected that PDB2PQR and Meeko were unavailable in its active environment, it seamlessly adapted, autonomously re-routing its pipeline to utilize OpenBabel and RDKit to add polar hydrogens and generate the required 3D conformers.

Following automated active-site identification via fpocket, flexible-ligand rigid-receptor docking with AutoDock Vina^30^ yielded a top-ranked binding affinity of -11.8 kcal/mol (**Figure 6A**). Imatinib inhibits Abl protein tyrosine kinase with an in vitro IC50 of 25-38 nM^31^, corresponding to an estimated binding free energy of -10.2 to -10.4 kcal/mol (ΔG = RT ln IC50, 298 K). The close agreement between the predicted and experimentally derived values supports the validity of the docking protocol. Pipette then autonomously superimposed the top docked conformation onto the co-crystallized reference ligand (**Figure 6B**). Notably, during this validation step, the agent encountered a common bioinformatics tool failure: a PDBQT-to-PDB conversion bug that stripped the molecule down to only three atoms. The agent autonomously diagnosed the parsing error, wrote custom code to extract coordinates directly from the PDBQT file, and recognized that standard RDKit RMSD calculations were artificially inflated (4.70 Å) due to internal atom-ordering ambiguities. To resolve this, the agent autonomously deployed a Hungarian optimal atom-matching algorithm. The resulting true heavy-atom RMSD was 0.89 Å with a centroid displacement of 0.43 Å, clearing the 2.0 Å threshold for successful pose reproduction^30^. Per-atom displacements are narrowly distributed, with the majority of heavy atoms displaced by less than 1.0 A from their crystallographic positions (**Figure 6C**). A summary of key validation metrics is shown in **Figure 6D**.

**Figure 6.**
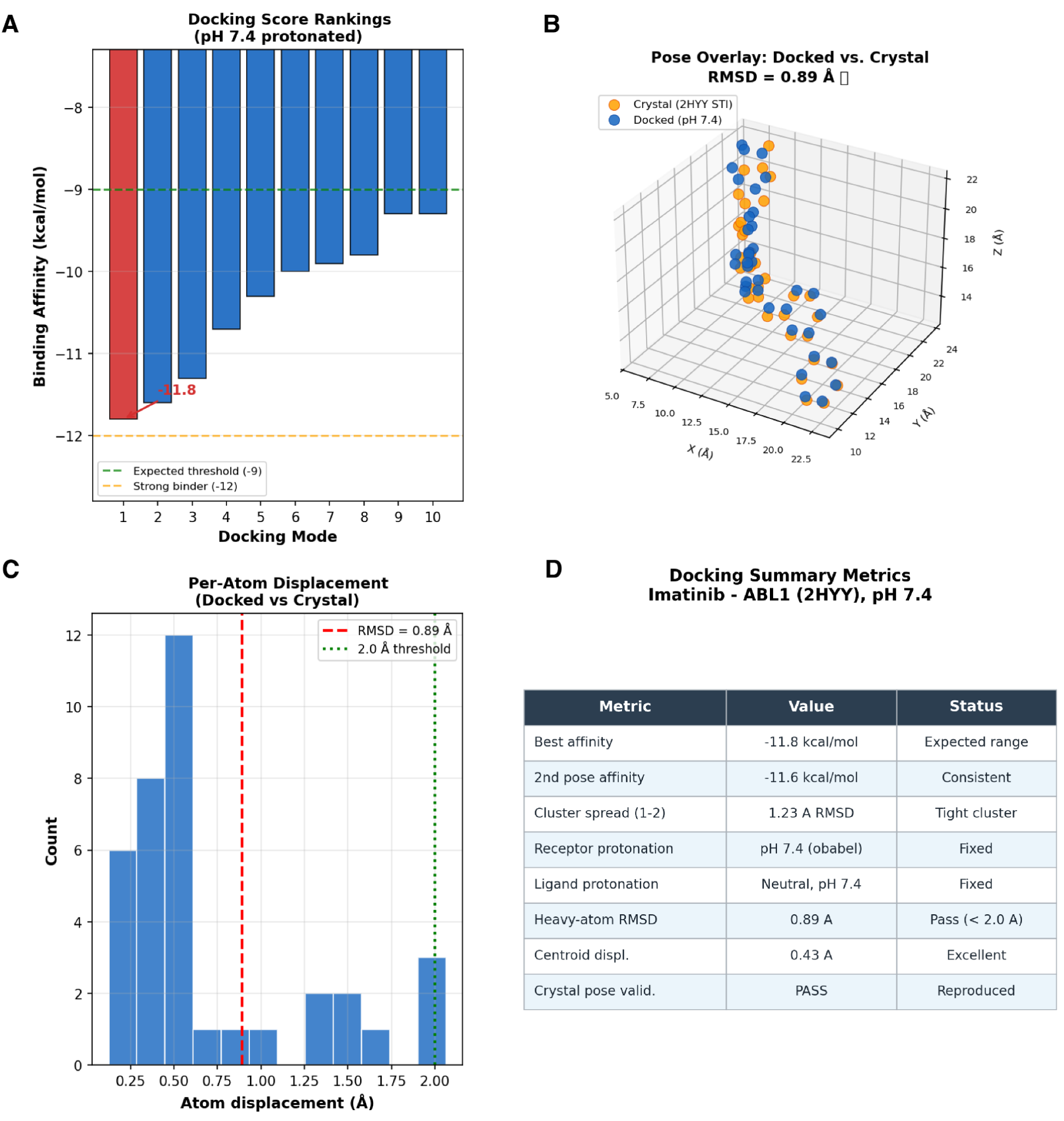
Molecular docking validation of imatinib into ABL1 kinase (PDB: 2HYY). **A)** Binding affinity rankings for the top 10 docking modes at pH 7.4 protonation state. The best-scoring pose achieved -11.8 kcal/mol (red), within the expected range from published crystal structure data. The dashed green line indicates the expected threshold (-8 kcal/mol) and the orange line marks the strong binder cutoff (-12 kcal/mol). **B)** Three-dimensional overlay of the docked pose (blue, pH 7.4) and the co-crystallized imatinib conformation (orange, PDB 2HYY), with a heavy-atom RMSD of 0.89 A. **(C)** Histogram of per-atom displacements between the docked and crystal poses. The majority of heavy atoms fall within 0.5-1.0 A; the red dashed line denotes the 2.0 A acceptance threshold commonly used in docking validation studies. **D)** Summary metrics table reporting best binding affinity (-11.8 kcal/mol), cluster spread (< 2.4 A RMSD between top poses), receptor and ligand protonation states (pH 7.4), heavy-atom RMSD (0.89 A), and crystal pose validation status. Docking was performed with AutoDock Vina against the ABL1 kinase domain; the sub-2.0 A RMSD confirms successful reproduction of the experimentally determined binding mode.

Beyond geometric precision, Pipette successfully recapitulated the pharmacophoric interaction network that defines imatinib’s clinical mechanism of action. The autonomous workflow correctly Å). The Asp381 interaction is consistent with imatinib’s classification as a type II inhibitor that selectively targets the inactive, DFG-out kinase conformation^32^. The Thr315 contact is directly relevant to the clinically significant T315I resistance mutation in chronic myeloid leukemia, where substitution by isoleucine abolishes this hydrogen bond and sterically occludes drug binding^32,33^

Thus, by autonomously navigating missing dependencies, bypassing software parsing bugs, deploying advanced symmetry-corrected algorithms, and ultimately generating sub-Angstrom structural predictions, Pipette demonstrates the capacity to execute highly resilient, clinically relevant computational chemistry workflows.

#### Drug Designing task 2: de novo cyclic peptide design

In the next task, we extended the drug design evaluation beyond small molecule docking. We tasked Pipette with de novo cyclic peptide design targeting the p53-MDM2 protein-protein interaction (PPI), a therapeutically important but structurally challenging target class. Given only the MDM2-p53 co-crystal structure (PDB 1YCR) and the instruction to design cyclic peptides mimicking the p53 hotspot triad (Phe19, Trp23, Leu26), the agent autonomously designed 10 head-to-tail lactam-cyclised peptides of 6-8 residues, each preserving the Phe-Trp-Leu pharmacophore at positions 1-3 while varying the remaining positions to explore hydrophobic fillers, polar contacts, charged rim interactions, and backbone rigidification strategies (**Figure 7A**). All 10 peptides were docked into the MDM2 hydrophobic cleft using AutoDock Vina, with two peptides (CP02, CP09) rescued by SMINA after Vina convergence failure due to excessive conformational flexibility. Binding affinities ranged from -7.4 to -12.1 kcal/mol (**Figure 7A**), with the top candidate CP07 (Tyr-Trp-Leu-Ala-Phe-Ala-Ala, 7-mer) achieving -12.1 kcal/mol – substantially exceeding the re-scored native p53 transactivation domain peptide (-6.8 kcal/mol on the same receptor).

**Figure 7.**
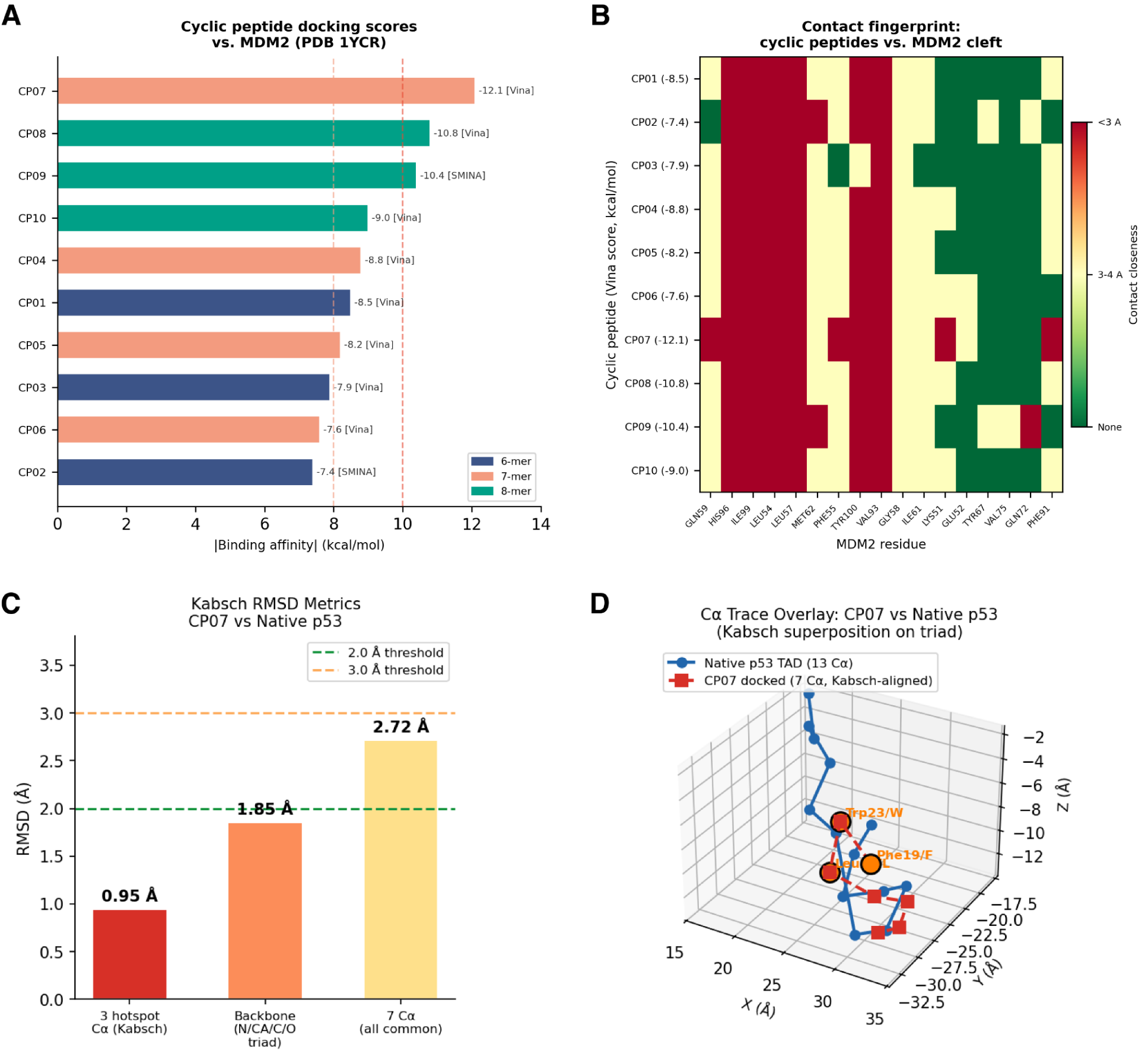
De novo cyclic peptide design targeting the p53-MDM2 interaction. **(A)** Docking scores for 10 cyclic peptides (6-8 residues) designed to mimic the p53 Phe19-Trp23-Leu26 hotspot triad, docked into the MDM2 hydrophobic cleft (PDB 1YCR). Bars are coloured by peptide length; docking engine indicated (Vina or SMINA rescue). Dashed lines mark score thresholds: red, excellent (< -10 kcal/mol); orange, strong (-8 to -10); green, moderate (-6 to -8). **(B)** Contact fingerprint heatmap showing minimum heavy-atom distance (A) between each peptide and MDM2 binding-cleft residues. Dark colours indicate close contact (< 3.0 A). All 10 peptides engage the canonical hotspot residues Ile99, Leu54, Leu57, His96, Val93, and Tyr100. **(C-E)** Structural comparison of the top-ranked peptide CP07 against the native p53 transactivation domain. **(C)** RMSD metrics at three levels of granularity (hotspot triad: 0.95 A; backbone triad: 1.85 A; all common C-alpha: 2.72 A). **(D)** C-alpha trace overlay after Kabsch superposition.

A contact fingerprint analysis confirmed that all 10 peptides engaged the canonical MDM2 hotspot residues (Ile99, Leu54, Leu57, His96, and Val93) with 100% occupancy, and Tyr100 was contacted by 9 of 10 designs (**Figure 7B**). Kabsch RMSD superposition of the top-ranked CP07 pose against the co-crystallised p53 peptide yielded a hotspot triad C-alpha RMSD of 0.95 Å, confirming sub-angstrom pharmacophore reproduction despite the fundamentally different backbone topology (cyclic vs. linear alpha-helix) (**Figure 7C**). The backbone triad RMSD (1.85 Å) and all-common-C-alpha RMSD (2.72 Å) reflect the expected topological divergence between cyclic and helical scaffolds. The Kabsch-aligned C-alpha trace overlay confirms spatial convergence of the hotspot triad between CP07 and native p53 (**Figure 7D**).

The Reviewer Agent flagged two failures during this analysis: incomplete docking (CP02 and CP09 initially lacked poses) and the use of centroid displacement instead of proper atom-level RMSD. The agent resolved both autonomously, rescuing the failed peptides with SMINA and implementing the Kabsch SVD superposition algorithm for RMSD validation. The reviewer additionally issued warnings about the inherent limitation of Vina for macrocyclic peptides (which are treated as branched flexible ligands rather than true rings) and the need for score calibration against a known reference, both of which were explicitly addressed in the final report.

By autonomously designing, building, docking, and structurally validating cyclic peptide candidates, and recovering from docking engine failures and incorrect RMSD methodology through the self-correction loop, Pipette demonstrates the capacity to execute de novo molecular design workflows that span medicinal chemistry, structural biology, and computational chemistry.

#### Clinical Variant analysis

To evaluate Pipette’s capacity for expert-level clinical reasoning, we tasked the agent with performing an ACMG Secondary Findings (v3.2)^34^ analysis on the Genome in a Bottle (GIAB) HG002 reference standard. Provided only with a raw, unannotated VCF and a plain-text list of the 81 ACMG gene symbols, Pipette autonomously orchestrated an end-to-end clinical diagnostic workflow. The agent successfully utilized its database-query skill to map the clinical panel to GRCh38 genomic coordinates, dynamically resolved a Java heap memory overflow during SnpEff annotation, and executed coordinate-based lookups against external databases including gnomAD and ClinVar.

Notably, Pipette demonstrated deep domain-specific clinical awareness that surpasses standard programmatic pipelines. The agent autonomously identified a critical discrepancy between the requested clinical assay and the input data, flagging the absence of Chromosome X in the input VCF and issuing a formal limitation warning that the X-linked genes GLA and OTC could not be evaluated.

To systematically process the genomic data, Pipette constructed and executed a stringent, multi-tiered variant filtering pipeline (**Figure 8A**). Starting with 9,924 high-quality (PASS filter) variants located within the evaluable ACMG gene boundaries, the agent performed functional annotation to isolate 101 candidate variants exhibiting HIGH or MODERATE functional impact, alongside relevant splice-region alterations. The final classification comprised 92 Benign, 9 VUS, and 0 Pathogenic or Likely Pathogenic variants (**Figure 8B**), consistent with the expected absence of reportable secondary findings in this healthy reference individual. VUS were distributed across seven genes (PRKAG2, HNF1A, DSG2, MSH2, NF2, TSC1, TTN), with the majority representing splice-region variants of low predicted impact or missense variants with conflicting ClinVar submissions (**Supplementary figure S2A**). Analysis of these 101 clinically relevant candidates revealed a characteristic distribution of molecular consequences, predominantly comprising missense variants (n=68) and splice region variants (n=31), with rare frameshift and disruptive in-frame deletions (**Supplementary figure S2B**).

**Figure 8.**
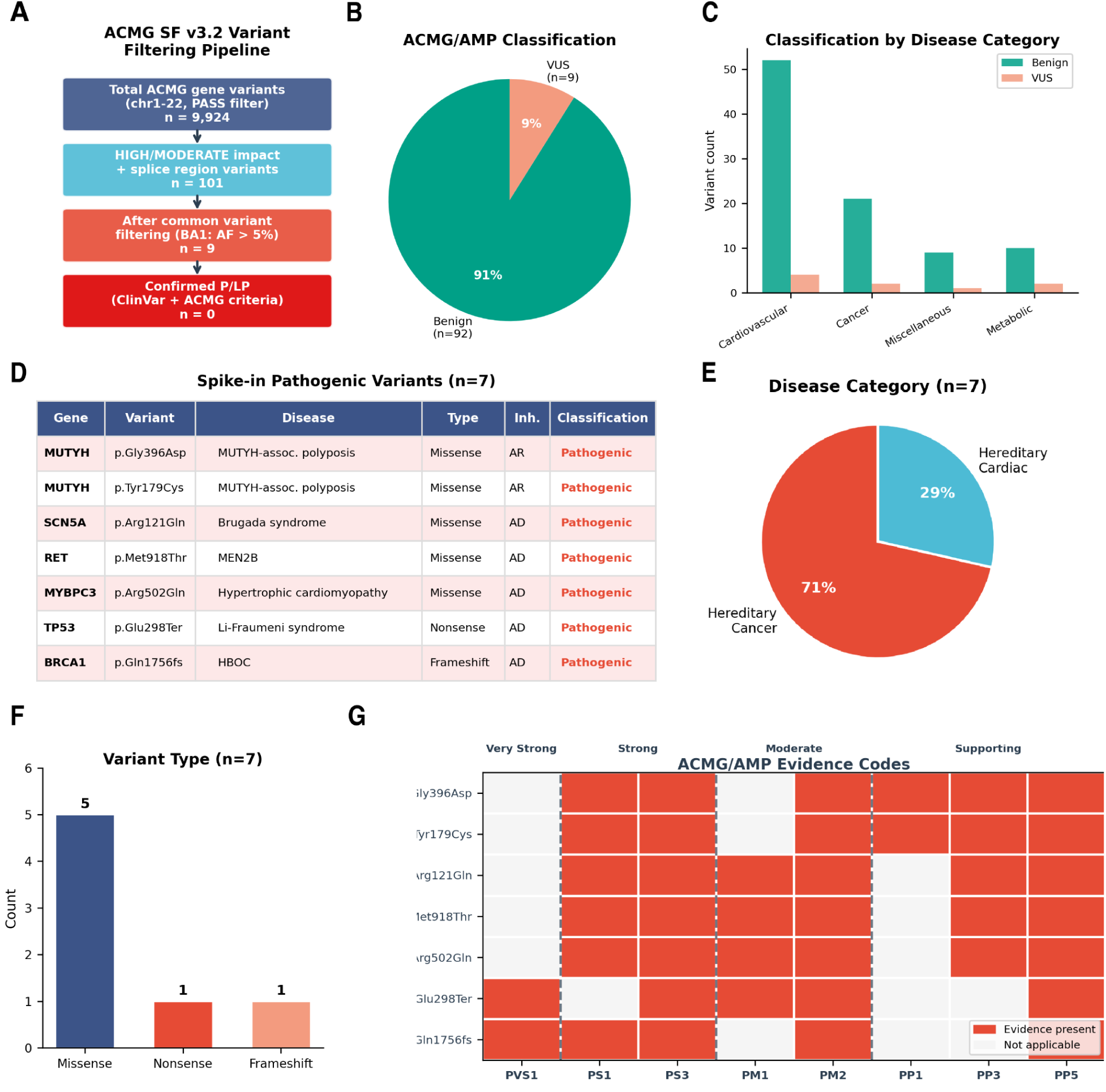
ACMG Secondary Findings v3.2 variant filtering and classification in HG002. **A)** Variant filtering pipeline showing progressive reduction from 9,924 PASS variants in ACMG gene regions (chr1-22) to 101 HIGH/MODERATE impact and splice region candidates, 9 variants remaining after common variant filtering (BA1: allele frequency > 5%), and 0 confirmed Pathogenic or Likely Pathogenic variants. **B)** ACMG/AMP classification of the 101 candidate variants: 92 (91%) classified as Benign/Likely Benign and 9 (9%) as Variants of Uncertain Significance (VUS). **C)** Classification breakdown by ACMG disease category, showing Cardiovascular genes contribute the most variants (Benign and VUS), followed by Cancer and Miscellaneous categories. **D)** Seven known pathogenic variants spanning six ACMG SF v3.2 genes were injected into the GIAB HG002 benchmark VCF (4,048,342 background variants) and analysed by Pipette. All seven were correctly classified as Pathogenic. The two MUTYH variants (autosomal recessive) were recognised as compound heterozygous biallelic, meeting the ACMG SF v3.2 reporting threshold for AR genes. No false-positive P/LP calls were generated from the background variants (specificity = 100%). AD, autosomal dominant; AR, autosomal recessive; HBOC, hereditary breast and ovarian cancer; MEN2B, multiple endocrine neoplasia type 2B. **E)** Distribution by variant consequence type and **F)** variant type. **G)** ACMG/AMP evidence code heatmap. Red cells indicate criteria met; grey cells indicate criteria not applicable. Codes are grouped by evidence strength: Very Strong (PVS1), Strong (PS1, PS3), Moderate (PM1, PM2), and Supporting (PP1, PP3, PP5). PVS1 (null variant) was applied only to the two loss-of-function variants (TP53 nonsense, BRCA1 frameshift). All variants received PS1 (same amino acid change as established pathogenic variant) and PS3 (functional studies demonstrating damaging effect). Classification rules followed Richards et al. (2015).

Pipette then autonomously queried population-level allele frequencies via the gnomAD v4 API to apply ACMG/AMP frequency criteria^35^ (**Supplementary figure S2C**). By strictly applying the BA1 criterion (allele frequency > 5% as a stand-alone benign assertion) and cross-referencing exact-allele matches in ClinVar, the agent seamlessly filtered out common population variation. Furthermore, the agent correctly avoided a common annotation artifact by demoting a spurious PRKAG2 frameshift – annotated by default against a non-canonical transcript – to Benign status due to its >5% allele frequency in the canonical transcript. The agent stratified the evaluated variants across all major ACMG disease categories, with the highest burden found in cardiovascular and cancer-associated genes (**Figure 8C**).

Further, we constructed a spike-in benchmark by injecting seven known pathogenic variants into the GIAB HG002 whole-genome VCF (GRCh38, chr1-22; see Methods). The spike-in set was designed to test sensitivity across three variant types (missense, nonsense, and frameshift), two inheritance modes (autosomal dominant and autosomal recessive), and five disease categories spanning oncology and cardiology. The set included a compound heterozygous MUTYH pair (c.1187G>A / p.Gly396Asp and c.536A>G / p.Tyr179Cys), which is only reportable under ACMG SF v3.2 when both alleles are pathogenic, essentially testing whether the agent correctly applies autosomal recessive reporting logic.

Pipette identified all seven spike-in variants (7/7 sensitivity) spanning five hereditary cancer genes (71%) and two hereditary cardiac genes (29%)(**Figure 8D,E**), with missense variants comprising the majority (5/7) alongside one nonsense and one frameshift variant and classified each as Pathogenic using appropriate ACMG/AMP evidence codes (**Figure 8F**). For the two loss-of-function variants – BRCA1 c.5266dupC (frameshift) and TP53 c.892G>T (nonsense) – the agent correctly applied PVS1 (null variant) as very strong evidence, combined with PS1, PS3, and PM2 (**Figure 8G**). For the five missense variants (MYBPC3, SCN5A, RET, and both MUTYH), the agent applied PS1 (same amino acid change as established pathogenic variant) and PS3 (functional studies), supplemented by PM1 (mutational hotspot), PM2 (population rarity), and PP3 (in silico predictions). No false-positive Pathogenic or Likely Pathogenic calls were generated from the ∼4 million background HG002 variants, yielding 100% specificity within the ACMG SF v3.2 gene panel.

The MUTYH compound heterozygosity test was handled correctly. The agent recognized that MUTYH is autosomal recessive in the ACMG SF v3.2 panel, identified both heterozygous variants, inferred biallelic status, and flagged the finding as reportable with a recommendation for parental phasing to confirm trans configuration. The agent also generated clinically appropriate per-variant management recommendations, including prophylactic thyroidectomy for RET p.Met918Thr (MEN2B), whole-body MRI surveillance for TP53 (Li-Fraumeni syndrome), and PARP inhibitor eligibility for BRCA1 carriers. Furthermore, Pipette triaged the seven pathogenic findings by clinical urgency, ranking RET p.Met918Thr (MEN2B) as urgent given the recommendation for prophylactic thyroidectomy within the first six months of life, followed by three high-priority findings requiring immediate specialist referral (TP53, BRCA1, SCN5A), and two moderate-priority findings warranting surveillance protocols (MYBPC3, MUTYH biallelic) **(Supplementary figure S2D).**

Together, these results demonstrate that Pipette can autonomously navigate complex genomic data to produce a clinically sound, highly interpretable diagnostic report that strictly adheres to established medical guidelines.

### Skill Graph-guided analysis achieves higher concordance with published results than unguided LLMs

To evaluate the contribution of the Skill Graph to analytical quality, we repeated three benchmark tasks – bulk RNA-seq differential expression, PBMC single-cell analysis, and human pancreas single-cell analysis – with the Skill Graph removed from Pipette’s knowledge base.

Because Pipette uses Claude Opus 4.5 as the main LLM, and GPT 5.4 as the fallback LLM, we performed these tests independently on both, given identical prompts and input data without access to the Skill Graph.

For the PBMC 68K single-cell benchmark, Claude subsampled the dataset, analyzed 20,000 of 68,579 cells (29%) and used manual marker scoring rather than automated annotation. GPT processed all cells but used MiniBatchKMeans (k=10) instead of graph-based clustering, and missed DC/pDC and megakaryocyte populations entirely **(Supplementary Figure 3A)**. Pipette (with Skill Graph) achieved the highest monocyte marker recall (86%) compared to Claude (29%) and GPT (43%), reflecting its Skill Graph-guided selection of CellTypist with the Immune_All_Low model (**Supplementary Figure 3B**). Across the seven matched cell types, Pipette achieved the lowest mean absolute error (4.3) relative to Zheng et al. 2017 reference proportions, compared with Claude 4.5 (4.4) and GPT 5.4 (4.5) (**Supplementary Figure 3C).**

For the human pancreas benchmark (GSE85241), all three systems correctly identified the major endocrine and exocrine cell types with similar proportional concordance (**Supplementary Figure 3D**). Pipette outperformed other two systems in marker gene recall among differentially expressed genes, with mean recall of 98% compared to Claude’s 87% and GPT’s 96% for the seven shared cell types (**Supplementary Figure 3E**). However, the systems differed in batch correction strategy; Pipette used scVI^36^, GPT used Harmony^37^, while Claude applied none. The systems also used a different clustering approach (**Supplementary Figure 3F**).

The three systems diverged most substantially in the rice leaf RNA-seq benchmark (GSE295637). GPT detected significantly fewer DEGs than published values at matched thresholds (**Supplementary Figure 4A**). Across all four contrasts, Pipette (with Skill Graph) achieved the highest Pearson correlations with the published log₂ fold changes (r = 0.976–0.991), followed by Claude (r = 0.929–0.980) and GPT (r = 0.914–0.976) (**Supplementary Figure 4B**). In terms of methodology, Pipette selected DESeq2 in R with a batch- and segment-corrected design formula (∼batch + segment + condition), matching the original publication. Claude used PyDESeq2 with a reduced ∼condition formula, omitting both covariates. GPT did not [perform any negative binomial GLM framework and instead performed ordinary least squares regression on log-transformed normalised counts (**Supplementary Figure 4C**).

Further, to test the Skill Graph’s ability to guide multi-step workflows spanning different analytical domains, we designed a cross-domain transition benchmark requiring three sequential skills: differential expression analysis, pathway enrichment, and drug target identification. Starting from an edgeR output table of 13,990 human genes (synthetic data), each system was prompted to select the top 500 upregulated DEGs, perform KEGG pathway enrichment, identify druggable targets, and query the OpenTargets platform for known compounds.

Pipette (with Skill Graph) correctly identified that the top 500 genes by fold change were dominated by non-coding RNAs (474/500), filtered to the 23 protein-coding genes, and applied clusterProfiler with a matched gene universe, yielding 3 statistically significant KEGG pathways (padj <= 0.05), all immune-related. It then queried the OpenTargets API for all 23 genes, identifying 8 druggable targets with 24 drug compounds, including 4 with FDA-approved drugs (CTLA4, KCNA3, CD52, CCR4). Claude Opus 4.5 also recognised the non-coding RNA issue and completed all three analysis steps, identifying 9 druggable genes and 12 drug compounds. However, its KEGG enrichment yielded no statistically significant pathways after multiple testing correction (lowest padj = 0.51). GPT 5.4 recognised the non-coding RNA composition but did not execute formal pathway enrichment or query the OpenTargets API, instead providing a qualitative interpretation of ∼2 candidate targets from its own internal knowledge. Seven of 10 druggable genes and 7 of 29 drug compounds were independently identified by both Pipette and Claude (**Supplementary Figure 5A,B**).

## Discussion

In this study, we presented Pipette, a multi-agent AI framework that translates natural language queries into reproducible bioinformatics workflows. By benchmarking the system across single-cell transcriptomics, bulk RNA-seq analysis, structure-based drug design, and clinical variant classification, we demonstrated that a literature-grounded agentic system can recapitulate established biological and clinical findings without requiring the user to write code or manage computational infrastructure.

A central design principle of Pipette is the separation of workflow planning from unconstrained code generation. Base LLMs are proficient at writing isolated scripts but are susceptible to hallucination^20^ and compounding methodological errors when orchestrating multi-step pipelines. The Skill Graph addresses this by constraining the agent’s planning space to analytical transitions that are supported by literature co-occurrence and validated by data type compatibility. The central methodological contribution, however, is not improved NER but the finding that document-level positional ordering validated against EDAM-derived data type constraints recovers an order of magnitude more pipeline connections (60.5% vs. 6.2% recall) than sentence-level RE. This aligns with observations from SciREX^25^ that document-level information extraction remains substantially harder than sentence-level tasks, but suggests that for workflow extraction specifically, the chronological structure of methods sections provides a strong inductive bias that supervised RE models cannot exploit.

The independent Reviewer Agent provides an additional layer of quality control through automated methodological auditing. The value of this self-correcting architecture was evident across our benchmarks: the agent autonomously recognized and remediated missing physiological pH protonation in the ABL1 kinase docking workflow, flagged the absence of chromosome X in a clinical reference VCF, and correctly demoted a spurious PRKAG2 variant initially misclassified as Likely Pathogenic due to a non-canonical transcript annotation artifact.

Reproducibility is a persistent challenge in computational biology, and Pipette’s provenance tracking system addresses this directly. Each executed workflow generates a machine-readable data lineage capturing software versions, random seeds, environmental configurations, and the full computational directed acyclic graph (DAG). In regulated domains such as clinical variant classification, where the agent documented its application of specific ACMG/AMP criteria against population databases, this level of analytical transparency is not a convenience but a requirement.

Several limitations should be noted. First, Pipette’s analytical repertoire is bounded by the Skill Graph. While the current graph covers the majority of standard bioinformatics workflows, novel algorithmic approaches that have not yet established a co-occurrence footprint in the published literature cannot be inferred. The graph will require periodic updating as methods evolve, and the mechanism for incorporating new tools and transitions at scale remains an area of active development. Second, all benchmarks in this study used published datasets with known biological outcomes. This enables direct comparison with established findings but does not constitute a prospective evaluation of the system on novel, unpublished data where ground truth is unavailable. Third, the automated literature retrieval module consistently favored recent keyword-matched publications over foundational references; in the docking benchmark, for example, the agent cited recent methodological papers but failed to retrieve the canonical crystal structure and pharmacological studies that directly underpin the analysis^31,32,38^. Fourth, while the Reviewer Agent caught several meaningful errors across our benchmarks, we did not systematically quantify its false negative rate (errors that pass review) or false positive rate (correct steps flagged as errors). A formal evaluation of reviewer accuracy across workflow types would strengthen confidence in this component. Fifth, the system depends on a commercial large language model, introducing vendor dependency and associated cost and latency considerations that we have not formally characterized. Finally, while the agent demonstrated standards-compliant application of ACMG/AMP classification guidelines, its outputs in clinical contexts must remain strictly investigational. All AI-generated clinical variant classifications require review and sign-off by a board-certified molecular geneticist before informing patient care.

Also to be noted is that the Skill Graph is constructed from a static corpus of approximately 20,000 papers and therefore represents a snapshot of the bioinformatics landscape at the time of extraction. As tools are deprecated and new methods emerge, the graph risks guiding users toward outdated workflows. We mitigate this in several ways. First, each skill is defined by an independently updatable specification (SKILL.md) that encodes explicit tool preferences and contraindications (e.g., prohibiting BWA for nanopore alignment), allowing curation without retraining the underlying NER/RE models. Second, edge weights reflect publication frequency, providing an implicit recency signal as newer methods accumulate citations. Third, the extraction pipeline is fully automated, enabling periodic corpus refresh and graph reconstruction. Future work will explore temporal weighting schemes that prioritize recent publications, automated detection of tool deprecation through version tracking and repository activity, and continuous graph updates triggered by new PubMed Central deposits.

Looking ahead, several extensions are planned. Implementing an automated Skill Graph update pipeline that continuously ingests new publications would reduce the manual curation burden and keep pace with rapidly evolving toolchains. Integrating generative molecular design tools would enable closed-loop drug discovery workflows from target identification through lead optimization. Adding GPU-accelerated compute environments would unlock access to biological foundation models for protein structure prediction, molecular generation, and single-cell embedding, substantially expanding the analytical capabilities available to the agent. Expanding support for multi-modal analyses that combine existing domains, such as linking single-cell transcriptomics with structure-based drug design in a single workflow, would better reflect the integrative nature of contemporary biological research. Finally, coupling Pipette’s computational outputs with autonomous robotic laboratory platforms would enable direct experimental validation of AI-generated hypotheses, closing the loop between in silico prediction and wet-lab confirmation.

By reducing the computational expertise required to execute standard genomic analyses, Pipette represents a step toward closing the gap between sequencing data generation and biological interpretation. As multi-omics datasets continue to grow in scale and complexity, literature-grounded, self-correcting agentic frameworks offer a promising approach to ensuring that analytical capacity keeps pace with data generation.

## Methods

### Construction of the Skill Graph

#### Data collection

We harvested ∼20,000 full-text papers from PubMed Central using NCBI E-utilities, sampling across 18 bioinformatics domains to ensure coverage of diverse tool ecosystems.

Domain-specific search queries targeted RNA-seq, variant calling, ChIP-seq, single-cell analysis, metagenomics, genome assembly, epigenomics, proteomics, long-read sequencing, metabolic flux analysis, clinical variants, pathogenomics, microbiome analysis, network analysis, drug design, CRISPR screening, GWAS, and next-generation sequencing. Methods sections were extracted from JATS XML using standard section heading detection. To supplement literature with explicit tool input/output descriptions, we also scraped ∼300 bioinformatics tutorials from Galaxy Training Network, Harvard Bioinformatics Core, nf-core pipeline documentation, Bioconductor package vignettes, and bioinformatics blog posts.

#### LLM-assisted annotation

We used GPT-4o for joint NER and relation extraction annotation. Each document was processed sentence by sentence with a structured prompt requesting entity spans (character offsets) and typed relations. A 50-document pilot with two-pass annotation verified prompt reliability before scaling to 800 documents. Systematic annotation errors identified during the pilot (incorrect PRODUCES examples, tool-subcommand splitting, verb-form operations) were corrected through prompt refinement.

#### Model training

Named entity recognition used PubMedBERT^39^ fine-tuned for token classification with IOB2 tags across three entity types. Training used 10 epochs, batch size 16, and learning rate 3e-5.

Relation extraction used PubMedBERT as a cross-encoder with typed entity markers. Input sentences were formatted with markers encoding entity type information. Training used focal loss (gamma=2.0) with 20 epochs and early stopping. A sentence-order constraint enforced that FOLLOWED_BY predictions required the head entity to appear before the tail entity in text.

Distant supervision from a curated gold knowledge graph provided additional training examples, weighted at 0.25x relative to manually annotated examples in the loss function.

The 800 annotated documents were split at the document level into training (560 documents, 70%), validation (120 documents, 15%), and held-out test (120 documents, 15%) sets using a fixed random seed (seed=42) to ensure reproducibility. No sentences from the same document appeared across splits. At the sentence level, the test set comprised 3,268 sentences for NER evaluation and 1,225 sentences (filtered to those containing ≥2 entities) for RE evaluation. All performance metrics reported in Figure 2B are computed exclusively on the held-out test set.

#### Pipeline extraction

Tool detection used dictionary matching with 215 regex patterns mapping tool names to 78 skill categories. For each paper, we recorded the text position of each tool mentioned and constructed ordered tool sequences per document. Consecutive skill pairs within each paper were aggregated into weighted FOLLOWED_BY edges, where the weight reflected the number of papers supporting each transition.

#### Data type validation

We curated input/output data type maps for all 78 skills using EDAM-aligned type names. Candidate pipeline edges were validated by checking whether the output types of the source skill intersected with the input types of the target skill. Edges passing this filter were labeled “literature+type.” Additional edges were inferred between skill pairs whose types were compatible but which were not observed together in any paper, labeled “type-inferred.“

#### Database query edge detection

Analysis of unmatched ground truth edges revealed that positional ordering and type validation miss a distinct class of workflow connections: database query edges. These represent analysis-to-lookup relationships (e.g., variant annotation followed by ClinVar query) where no data type flows between steps. We detected these edges through co-occurrence: for each paper containing both an analysis skill and a database query skill, we inferred a directed edge based on domain-informed rules. Data source databases (GEO, Ensembl, UCSC) were assigned as upstream of analysis steps, while annotation databases (ClinVar, COSMIC, STRING, KEGG) were assigned as downstream. Query-to-query cross-reference edges (e.g., ClinVar to COSMIC) were included when both databases co-occurred in the same paper. A minimum support threshold of 2 papers was applied.

#### Graph curation

The initial knowledge graph (2,094 nodes, 1,913 edges) built from annotated data underwent programmatic cleanup (removal of generic operations and wet-lab terms, merging of duplicates) followed by independent review from two LLMs (Claude Sonnet and Gemini). Edges flagged as incorrect by either reviewer were removed (union consensus). The resulting gold graph (1,679 nodes, 1,543 edges) served as the knowledge base for distant supervision^40^ and tool-to-skill mapping.

### Benchmark Datasets

All analyses were executed through the Pipette.bio conversational interface using default parameter selection guided by the platform’s Skill Graph layer. Complete parameter records, session files, and agent reasoning for each analysis are provided in the github repository https://github.com/variomeanalytics/pipette_benchmark.

Key prompts and parameters for each case study are as follows:

#### Rice bulk RNA-seq

The agent was prompted the following:

> *“Get the raw count matrix for this GSE295637 and pull the count matrix. Fetch the metadata from the series matrix file and find DEGs for these four contrasts: D30 vs W30, W40 vs W30, D40 vs W30, D40 vs D30. Give me full list of genes, along with fold change values and p values”*

In response, the agent loaded the ‘*geo-query’* skill first and downloaded the count matrix and the corresponding metadata file from the given GEO accession. It then loaded the *‘differential-expression’* skill, which guided it to identify the experimental design from the metadata (4 conditions × 5 leaf segments × 4 biological replicates = 80 total samples) and implement the statistical model.

*DESeq*2 *formula*: ∼*batch* + *segment* + *condition*

- batch (1 or 2): Controls for the sequencing batch effect (Batch1: reps 1–2, Batch2: reps 3–4; ∼40% read depth difference between batches)
- segment (seg2–seg6): Controls for positional variation along the leaf
- condition : Factor of interest; reference level = W30

The agent generated output files, plots, and reasoning are documented in the session summary available as a PDF from the project Github repository (see Methods).

#### PBMC scRNA-seq

For the sRNA-seq dataset, we obtained the matrix.mtx, barcodes.tsv, genes.tsv files from 10x Genomics public dataset (Fresh 68K PBMCs from a healthy human donor). Pipette was then prompted

> *“I uploaded a single-cell RNA-seq dataset in a file called pbmc68k.zip. This is single-cell RNA-seq data of Peripheral Blood Mononuclear Cells. Perform standard QC, normalize, find highly variable genes, run PCA, compute neighbors, cluster cells and identify marker genes per cluster. Annotate cell types based on known markers. Generate UMAP plots colored by cluster and by cell type, a dotplot of marker genes, and export the DEG table per cluster as CSV.”*

In response, the agent used the SCANPY^41^ to load the dataset (`scanpy.read_10x_mtx()`) and computed QC metrics with `sc.pp.calculate_qc_metrics(),`including 13 mitochondrial genes (prefix MT- ). It then applied filters: `min_genes=200, max_genes<3000` (doublet guard), `pct_counts_mt<20% , min_cells=3.` The agent then performed library-size normalization to 10,000 counts/cell (`sc.pp.normalize_total`), performed Log1p transformation, and saved `.raw` saved prior to subsetting top 2000 highly variable genes HVG to preserve all 17,788 genes for DE. The agent then scaled the data using `sc.pp.scale()` (zero-centered) and used 30 PCs for neighbor graph `sc.pp.neighbors(n_neighbors=20, n_pcs=30)`to plot the UMAP. Next, the agent applied the Leiden algorithm (`resolution=0.5`) to yield 16 clusters. To annotate the clusters, it used CellTypist (Immune_All_Low.pkl model) and majority-voting smoothed per cluster. Differential expression was detected using Wilcoxon rank-sum test with Bonferroni correction (`sc.tl.rank_genes_groups` ) run on `.raw` (all 17,788 genes) and filtered for padj <= 0.05.

#### Pancreas scRNA-seq

For the pancreas scRNA-seq task, we obtained the published human pancreas CEL-seq2 dataset (GSE85241)^27^ from GEO and prompted the agent:

> “*I have uploaded a single-cell RNA-seq count matrix file GSE85241_cellsystems_dataset_4donors_updated.csv.gz from GEO. This is a human pancreas study (GSE85241, CEL-seq2 protocol, 4 donors). Perform a complete single-cell analysis: quality control filtering, normalization, highly variable gene selection, dimensionality reduction (PCA), neighborhood graph construction, clustering, UMAP visualization, cell type annotation, and differential expression analysis. Identify the major pancreatic cell types and provide marker genes for each cluster. Generate a final report with UMAP plots colored by cluster and cell type, a dotplot of top marker genes, and a summary table of cell type proportions*.”

The agent loaded the dataset (GSE85241; 19,140 genes x 3,072 cells across 4 donors) using Scanpy. Gene names were stripped of chromosome suffixes ( chrXX). As mitochondrial genes were absent from this nuclear-fraction dataset, quality control filtering relied on gene count thresholds: min_genes=500 per cell (removing empty/dead wells), max_genes=6,000 per cell (removing doublets), and min_cells=5 per gene, retaining 1,795 cells and 17,167 genes. The agent normalised to 10,000 counts per cell (`sc.pp.normalize_total`), applied log1p transformation, and saved the full gene set to .raw prior to subsetting. Two thousand highly variable genes were selected using the Seurat dispersion method^42^ with the donor ID as batch key. The agent computed 50 PCA components (arpack solver), constructed 20-nearest-neighbour graph on the top 30 PCs, and generated a UMAP embedding (min_dist=0.3, spread=1.0). Clustering was performed with the Leiden algorithm at resolution 0.5, yielding 10 clusters. Cell type annotation was performed by overlap-score matching of the top 50 differentially expressed genes per cluster (Wilcoxon rank-sum test with Bonferroni correction) against a curated pancreatic marker dictionary including canonical markers for alpha (GCG, IRX2, ARX), beta (INS, MAFA, HADH), delta (SST, HHEX, LEPR), PP (PPY, ETV1), acinar (CTRB2, PRSS1, REG1A), ductal (KRT19, CFTR, SLC4A4), and stellate/mesenchymal (COL4A1, SPARC) cell types. Differential expression for marker gene identification was run on the .raw layer (all 17,167 genes) using `sc.tl.rank_genes_groups` with Bonferroni correction at p <= 0.05.

#### Drug Designing task-1

For the drug design task-1, the agent was prompted

> *“Dock imatinib (Gleevec) against the ABL1 kinase crystal structure PDB ID 2HYY. Retrieve the protein structure from PDB, prepare it by water and co-crystallized ligands, detect binding sites with fpocket, prepare imatinib from its SMILES (CC1=C(C=C(C=C1)NC(=O)C2=CC=C(C=C2)CN3CCN(CC3)C)NC4=NC=CC(=N4)C5=CN=CC=C5), generate 3D conformers, and perform molecular docking with AutoDock Vina. Report the top binding pose energy (expected: -9 to -12 kcal/mol), binding interactions, and generate a visualization of the docked pose.”*

The agent retrieved the ABL1 kinase crystal structure (PDB: 2HYY, 2.10 A resolution, chain A) and removed all HETATM records (water molecules, co-crystallised ligands). It added polar hydrogens at physiological pH using OpenBabel v3.1.0^43^ (`obabel -p 7.4`), expanding the receptor from 2,073 to 4,026 atoms, then converted to PDBQT format with Gasteiger partial charges (`obabel -xr --partialcharge gasteiger)`. For the ligand, the agent generated imatinib from its canonical SMILES and performed pKa analysis of the three ionisable nitrogens: piperazinyl N1 (pKa ∼8.1, ∼80% neutral at pH 7.4), piperazinyl N4 (pKa ∼3.7, fully deprotonated), and pyridyl N (pKa ∼1.7, neutral), selecting the neutral species as dominant at physiological pH. A 3D conformer was generated using RDKit ETKDGv3 with MMFF94 force field optimisation (randomSeed=42), yielding 8 active rotatable bonds, and converted to PDBQT with pH 7.4 protonation (`obabel -p 7.4 --partialcharge gasteiger`). The agent identified the ATP-binding site using fpocket^44^ (`fpocket -f receptor_clean.pdb`); the top-ranked pocket centroid (14.25, 15.28, 17.63 A) matched the co-crystallised STI ligand position. Flexible-ligand rigid-receptor docking was performed with AutoDock Vina 1.1.2^30^ using a 28 x 28 x 28 A search box (`exhaustiveness=32, num_modes=10, energy_range=4`).

To validate pose accuracy, the agent extracted 37 heavy atoms from both the best docked pose and the crystallographic imatinib, computed a pairwise distance matrix, and applied the Hungarian optimal atom-matching algorithm via `scipy.optimize.linear_sum_assignment` to obtain the order-independent heavy-atom RMSD. The 2.0 A threshold was applied as the acceptance criterion for successful pose reproduction^30^.

#### Drug Designing task-2

For the drug design task-2, the agent was prompted

> *“Design a set of 10 cyclic peptides (6-8 residues) targeting the p53-MDM2 interaction interface. Use the co-crystal structure PDB 1YCR as the reference. The peptides should mimic the key hotspot residues of p53 (Phe19, Trp23, Leu26) that bind the MDM2 hydrophobic cleft. For each peptide, generate a 3D conformer, dock into the MDM2 binding cleft, and report binding affinity and key contacts. Compare the top-ranked peptide against the co-crystallised p53 peptide pose. Report RMSD, binding score, and interaction fingerprint”*

The agent retrieved the MDM2-p53 co-crystal structure^45^ (PDB: 1YCR) and extracted chain A (MDM2, residues 17-111), removing water molecules and non-MDM2 heteroatom entries.

Hydrogens were added at physiological pH using OpenBabel^43^ and the structure was converted to rigid-receptor PDBQT format. The docking search box was centred on the co-crystallised p53 peptide centroid (25.42, -25.14, -7.92 A) with 30 x 30 x 30 A dimensions to encompass the full hydrophobic cleft and rim contacts. The agent then designed 10 head-to-tail lactam-cyclised peptides of 6-8 residues, each preserving the Phe-Trp-Leu hotspot triad at positions 1-3, mimicking the three p53 residues (Phe19, Trp23, Leu26) that contribute >75% of the binding energy^45^, while varying the remaining positions to explore hydrophobic fillers (Ala, Val, Ile, Leu), polar contacts (Ser, Thr, Asn, Gln), charged rim interactions (Lys, Glu, Asp, Arg), aromatic stacking (Tyr, His), and backbone rigidification (Aib). Three-dimensional conformers were generated using RDKit (ETKDGv3, 200 attempts) with MMFF94 force field minimisation, and PDBQT files were prepared via Meeko. Flexible-ligand rigid-receptor docking was performed with AutoDock Vina 1.1.2 at exhaustiveness 32, generating 10 poses per peptide with an energy range of 5 kcal/mol. Two peptides (CP02, CP09) failed Vina convergence due to excessive conformational flexibility; the agent autonomously re-docked these with SMINA^46^. Contact analysis was performed using BioPython NeighborSearch^47^ at a 4.5 A heavy-atom cutoff, recording per-residue minimum distances. To validate the top-ranked pose (CP07) against the native p53 peptide geometry, the agent implemented the Kabsch SVD superposition algorithm^48^ on matched backbone atoms (N, C-alpha, C, O), computing three RMSD metrics: hotspot triad C-alpha (3 atoms), backbone triad (12 atoms), and all common C-alpha (7 atoms).

#### Clinical Variant analysis

To benchmark clinical variant analysis workflows, we obtained the HG002_GRCh38_1_22_v4.2.1_benchmark.vcf.gz file from the Genome In A Bottle Consortium website (https://ftp-trace.ncbi.nlm.nih.gov/ReferenceSamples/giab/release/AshkenazimTrio/HG002_NA24385_son/NISTv4.2.1/GRCh38/). The VCF, along with ACMG SF v3.2 gene list^34^, was provided to Pipette and instructed for analysis in the following prompt:

> *“I have uploaded a patient’s whole genome VCF file HG002_GRCh38_1_22_v4.2.1_benchmark.vcf.gz and its index (aligned to the GRCh38 reference build), along with a text file containing the ACMG Secondary Findings v3.2 panel in ACMG_SF_v3.2_panel. I need to know if this patient carries any actionable secondary findings. Please analyze the genome to identify any ‘Pathogenic’ or ‘Likely Pathogenic’ variants strictly within the genes listed in the provided ACMG panel. Evaluate the variants using standard ACMG/AMP clinical interpretation guidelines. Please provide a final report detailing any clinically significant variants you find, including the specific ACMG evidence criteria (e.g., PVS1, PM2, etc.) used to classify them.”*

The agent first loaded the ‘clinical-variant’ skill and used it to extract PASS-filtered variants within the 81 ACMG SF v3.2 gene bodies using bcftools^49^ (v1.17+) with a ±2 kb boundary, using gene coordinates fetched from the Ensembl REST API (GRCh38, release 112)^50^, yielding 10,212 variants. Functional annotation was performed with SnpEff v5.4^51^ using the GRCh38.99 database. Variants with HIGH or MODERATE predicted impact were retained (n=70), along with LOW-impact splice region variants (n=31), totalling 101 candidates. The agent then loaded the ‘*gnomad-query*’ skill following the Skill Graph path, and queried the population allele frequencies from gnomAD v4^52^ via the GraphQL API. It applied the ACMG/AMP benign criteria: BA1 (allele frequency >5%, stand-alone benign) and BS1 (>0.1% for autosomal dominant conditions). For pathogenicity assessment, it loaded the *‘clinvar-query’* skill to obtain the ClinVar VCF^53^ (GRCh38, March 2025 release) and executed coordinate-plus-allele matching (CHROM, POS, REF, ALT) against all 101 candidate variants; no coordinate-only matches were accepted. Final variant classification followed the ACMG/AMP 2015 guidelines^35^ with 2018 ClinGen specifications, evaluating criteria PVS1, PS1, PS2, PM1, PM2, PM4, PM5, PM6, PP2, PP3, BA1, BS1, and BS2 per variant. For the spike-in benchmark, seven ClinVar Pathogenic variants across six ACMG SF v3.2 genes were injected into the same HG002 VCF using bcftools concat, and the merged file was re-indexed with tabix prior to analysis.

## Data and Code Availability

All datasets used in this study are publicly available through the NCBI Sequence Read Archive and 10X Genomics. Complete conversation logs, provenance records, and output files for all seven case studies are available at https://github.com/variomeanalytics/pipette_benchmark.

Large data files (DESeq2 RDS object and spiked-in VCF) are deposited at Zenodo (10.5281/zenodo.19433635). Pipette.bio is accessible at https://pipette.bio and the skill graph is accessible at http://skillgraph.pipette.bio.

## Author Contributions

C.G. conceived the project, designed the multi-agent architecture, developed the Skill Graph NLP pipeline (NER/RE training, graph construction, and type inference), built the skill execution framework, and acquired funding. A.S. curated training data for the Skill Graph, designed and executed the benchmark evaluations, and validated agent outputs against published results. Both authors analyzed the results and wrote the manuscript.

## Acknowledgements

The authors would like to thank Anadi Vasta for helping us develop and secure the cloud compute infrastructure, and Gaurav Uppal for helping us develop the user interface. The authors would also like to thank Dr. Venkategowda Ramegowda and his lab members at GKVK campus, University of Agricultural Sciences, Bangalore for testing the RNA-seq workflows on independent datasets, and Dr. Daifeng Wang, University of Wisconsin, Madison for inputs on the platform design. The authors would also like to thank early testers who gave active feedback on the agent.

