## Supplementary Figures for "Pipette: Encoding scientific literature into an executable Skill Graph for multi-agent bioinformatics"

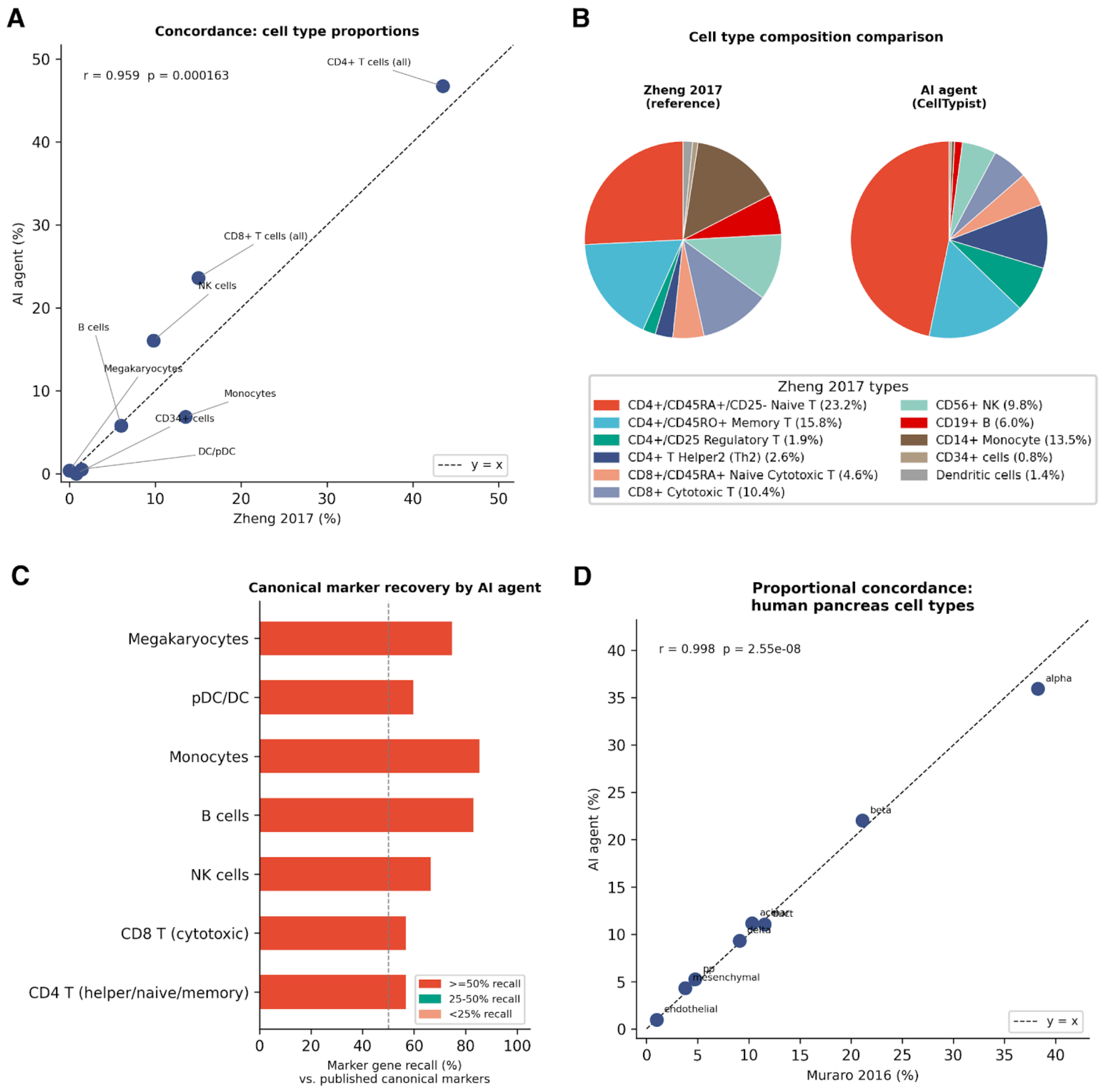


**Supplementary Figure S1**: **Evaluation of Pipette to execute scRNA-seq workflows using the PBMC 68K dataset. A)** Scatter plot of cell type percentages reported by the original study Zheng 2017 (x-axis) versus those identified by the Pipette (y-axis) for seven matched cell groups. Each point represents one cell type group. The dashed line indicates perfect concordance (y = x). Pearson correlation coefficient (r) and p-value are shown. **B)** Left: Cell type proportions reported by Zheng et al. 2017 using reference-based classification into 11 purified PBMC subpopulations. Right: Cell type proportions identified by Pipette using Leiden clustering and CellTypist annotation, resolved into 10 categories. The overall immune landscape is qualitatively concordant between the two methods: CD4+ T cells dominate in both (~44-47%), followed by CD8+ T cells (~15-24%), NK cells (~10-16%), monocytes (~7-14%), and B cells (~6%). Pipette additionally resolves classical versus non-classical monocyte subsets and CD56bright versus CD56dim NK populations not distinguished in the original publication. CD34+ progenitor cells (0.8% in Zheng 2017) are absent from the AI annotation, as the CellTypist `Immune_All_Low` model targets mature immune lineages only. **D)** Proportional concordance scatter plot. Each point represents one cell type; the dashed line indicates perfect agreement. Pearson r = 0.998 ($p = 2.55 \times 10^{-8}$).

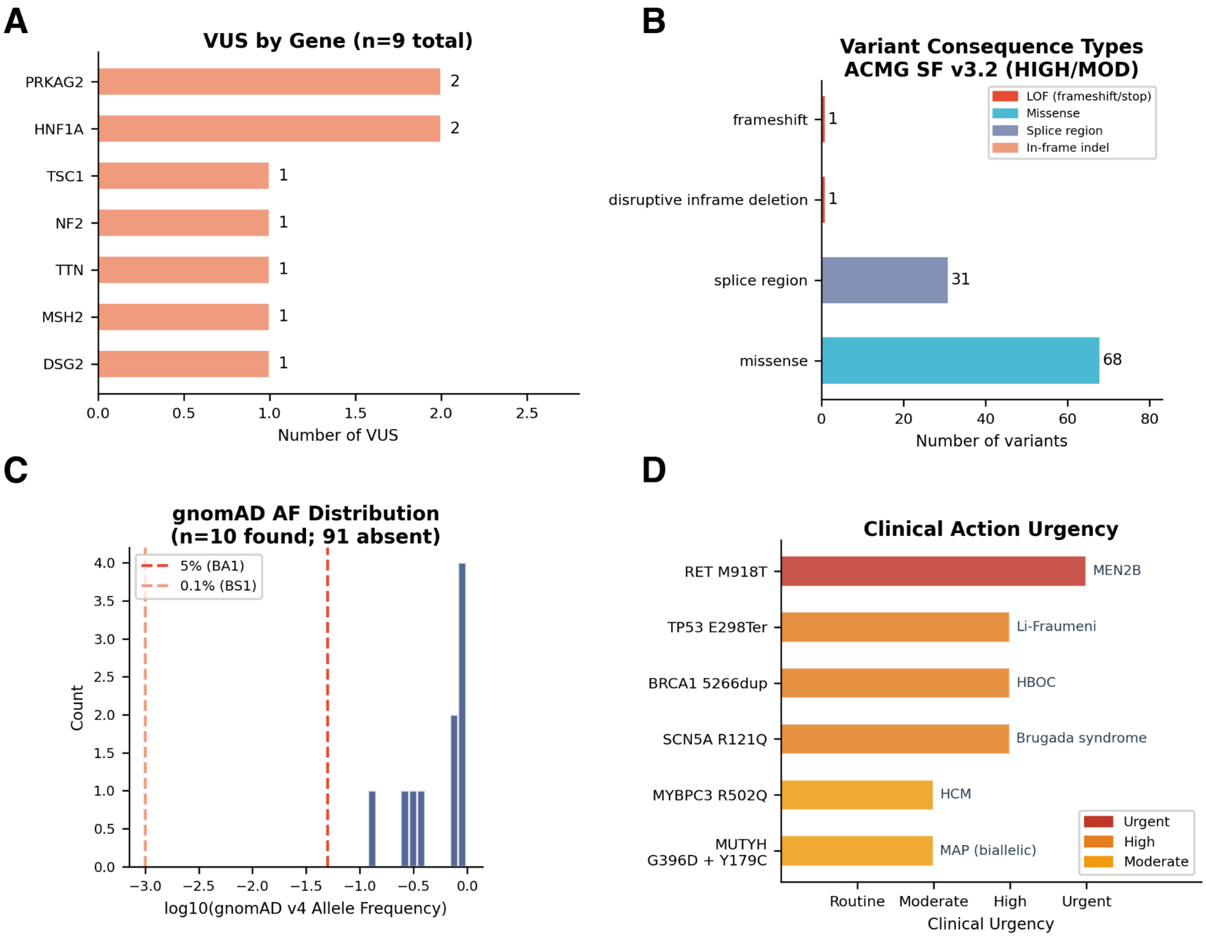


**Supplementary Figure S2: ACMG Secondary Findings v3.2 variant filtering and classification in HG002**. **A)** Barchart shows VUS distribution across 7 genes, with PRKAG2 and HNF1A each contributing 2 VUS. **B)** Variant consequence types among the 101 HIGH/MODERATE impact candidates in ACMG SF v3.2 genes: missense variants predominate (n=63), followed by splice region variants (n=31), with single frameshift and disruptive in-frame deletion events. **C)** Distribution of gnomAD v4 allele frequencies for the 10 variants with population data (91 absent from gnomAD); dashed lines indicate BA1 (5%) and BS1 (0.1%) benign evidence thresholds. **D)** Urgency tiers were determined by the agent based on time-sensitivity of recommended clinical interventions per ACMG/AMP guidelines and published management protocols. No standardised urgency scoring framework was applied; tiers reflect the agent's synthesis of clinical guideline recommendations from the corresponding analysis report.

**A**

**Methodology Comparison (PBMC 68K)**

| | Pipette | Claude 4.5 | GPT 5.4 |
|---|---|---|---|
| Cells analyzed | 68,548 (100%) | 20,000 (29%) | 68,263 (100%) |
| Tool | Scanpy | Scanpy | Scanpy (custom) |
| Clustering | Leiden (res=0.5) | Leiden (res=0.8) | KMeans (k=10) |
| Annotation | CellTypist (auto) | Marker scoring | Marker scoring |
| QC: max genes | 3,000 | 5,000 | 2,500 |
| QC: max MT% | 20% | 10% | 5% |
| Clusters | 16 | 13 | 10 |
| Cell types | 10 | 9 | 8 (no DC/Mega) |

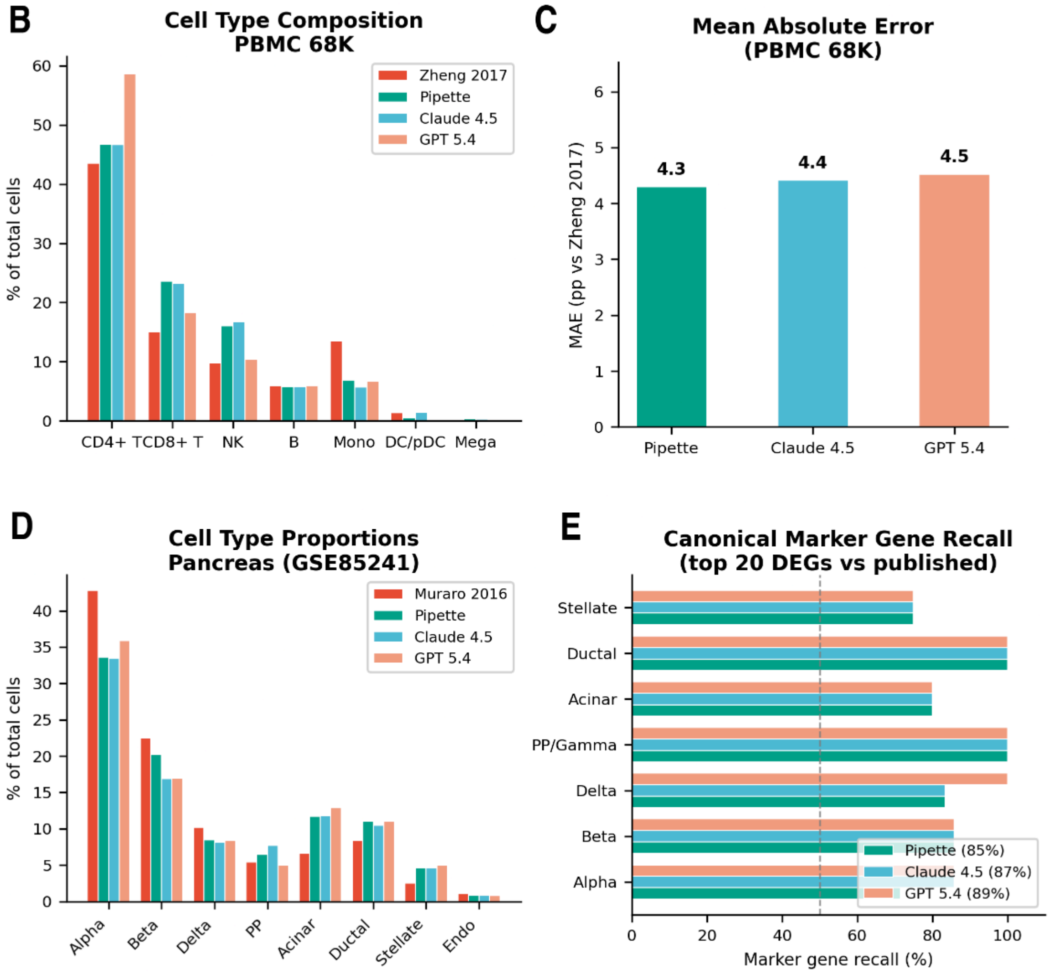


**F**

**Methodology Comparison (Pancreas)**

| | Pipette | Claude 4.5 | GPT 5.4 |
|---|---|---|---|
| Cells retained | 2,414 (79%) | 2,429 (79%) | 2,301 (75%) |
| Tool | Scanpy | Scanpy | Scanpy (custom) |
| Batch correction | scVI | None | Harmony |
| Clustering | Leiden (res=0.6) | Leiden (res=0.8) | Louvain (res=0.8) |
| Annotation | Marker dict (pancreas) | Marker dict | Marker dict |
| QC method | MAD-based | Hard cutoffs | Donor-aware MAD |
| Cell types | 10 | 11 | 9 |
| Endothelial | Yes (0.9%) | Yes (0.9%) | Yes (0.8%) |
| Macrophage | Yes (1.3%) | Yes (0.8%) | No |

**Supplementary Figure S3: Skill Graph ablation single-cell study. A)** A summary table of methodology comparison on the PBMC68K task with and without the Skill Graph (Claude Opus4.5 and GPT4.5). **B)** Grouped bar chart comparing the percentage of total cells assigned to each cell type by the three agentic systems and the original publication (Zheng et al. 2017). **C)** Mean absolute error (MAE) of cell-type proportion estimates relative to the Zheng et al. (2017) reference, computed across seven harmonized cell-type categories (CD4+ T, CD8+ T, NK, B, Monocytes, DC/pDC, Megakaryocytes). Each bar represents the average absolute deviation (in percentage) between an agent's reported proportions and the published reference values. **D)** Grouped bar chart comparing the percentage of total cells assigned to each cell type by the three agentic systems and the original publication of the pancreas dataset. **E)** Percentage of canonical marker genes recovered within the top 50 differentially expressed genes for each of seven pancreatic cell types. Recall was computed as the fraction of curated marker genes (from Muraro et al. 2016) found among the 50 highest-ranked DEGs per cell type in each agent's output. **F)** Summary table of methodology comparison on the pancreas task with and without the Skill Graph.

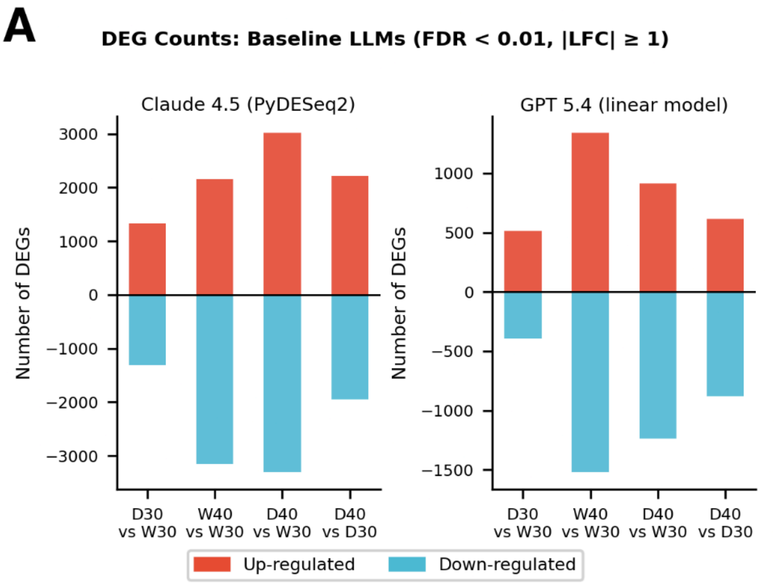


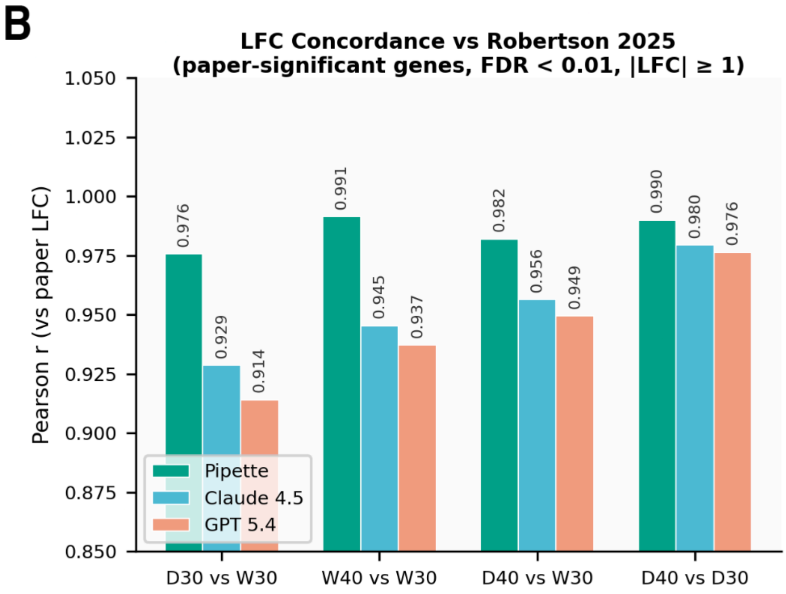


**C**

**RNA-seq Methodology Comparison**

| | Pipette | Claude 4.5 | GPT 5.4 |
|---|---|---|---|
| **Tool** | DESeq2 (R) | PyDESeq2 (Py) | Linear model (Py) |
| **Model** | NB GLM | NB GLM | OLS log2(n+0.5) |
| **Formula** | ~bat+seg+cond | ~condition | ~cond+segment |
| **Batch corr.** | Yes | No | No |
| **Seg. covar.** | Yes | No | Yes |
| **Pre-filter** | ≥10 in ≥4<br>24,338 | ≥10 in ≥5<br>24,049 | None<br>42,189 |

**Supplementary Figure S4: Skill Graph ablation bulk RNA-seq study. A)** Number of up-regulated (red) and down-regulated (blue) differentially expressed genes identified by Claude Opus 4.5 and GPT 5.4 (linear model, ~condition+segment) across four stress contrasts. **B)** Pearson correlation coefficients between each agent's log2 fold-change estimates and published values from Robertson et al. (2025), computed on paper-significant genes (FDR < 0.01, |LFC| ≥ 1). Pipette (green) achieved the highest concordance across all four contrasts (r = 0.976–0.991), followed by Claude 4.5 (blue; r = 0.929–0.980) and ChatGPT 5.4 (orange; r = 0.914–0.976). **C)** Summary of analytical choices made by each agent.

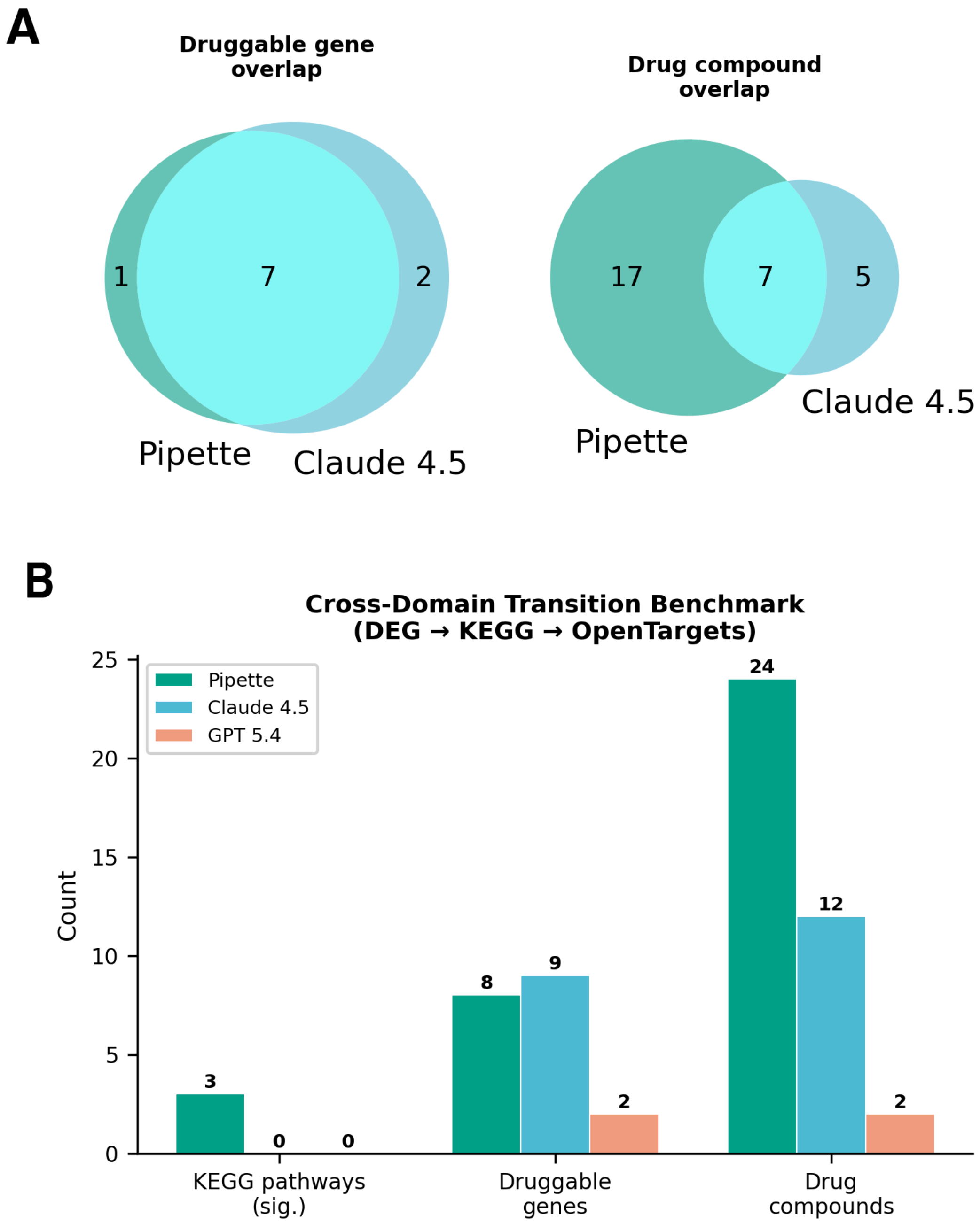


**Supplementary Figure S5: Skill Graph ablation study for multi-domain query. A)** Druggable gene and drug compound overlap between Pipette and Claude 4.5. Venn diagrams showing the intersection of druggable gene targets (left) and drug compounds (right) identified by Pipette and Claude 4.5 through the DEG-to-drug-target pipeline. **B)** Comparison of the number of significant KEGG pathways (padj < 0.05), druggable gene targets, and drug compounds identified by each agent across the full DEG → KEGG → OpenTargets pipeline.
